# A conjugation platform for CRISPR-Cas9 allows efficient β-cell engineering

**DOI:** 10.1101/732354

**Authors:** Donghyun Lim, Vedagopuram Sreekanth, Kurt J. Cox, Benjamin K. Law, Bridget K. Wagner, Jeffrey M. Karp, Amit Choudhary

## Abstract

Genetically fusing protein domains to Cas9 has yielded several transformative technologies; however, these fusions are polypeptidic, limited to the Cas9 termini and lack multivalent display, and exclude diverse array of molecules. Here, we report a platform for the site-specific and multivalent display of a wide assortment of molecules on both the termini and internal sites on Cas9. Using this platform, we endow Cas9 with the functionality to effect precision genome edits, which involves efficient incorporation of exogenously supplied single-stranded oligonucleotide donor (ssODN) at the break site. We demonstrate that the multivalent display of ssODN on Cas9 significantly increased precision genome edits over those of Cas9 bearing one or no ssODN, and such display platform is compatible with large oligonucleotides and rapid screening of ssODNs. By hijacking the insulin secretion machinery and leveraging the ssODN display platform, we successfully engineer pancreatic β cells to secrete protective immunomodulatory factor interleukin-10.

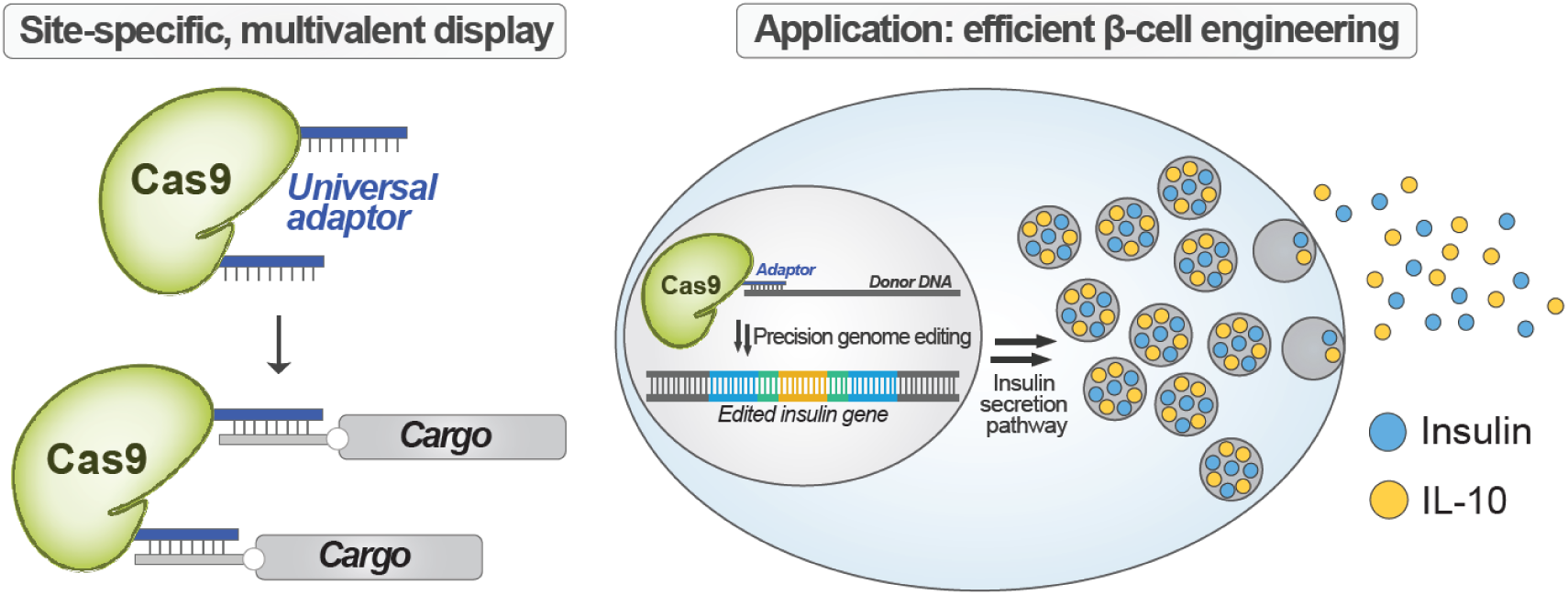
TOC GRAPHIC.

---

CRISPR-Cas9 is a DNA endonuclease that can be targeted to a genomic site using a guide RNA (gRNA) bearing sequence complementarity to the target site.^1^ Genetic fusion of Cas9 with effector domains (e.g., a transcription activator) has yielded transformative technologies;^2,3^ however, this approach is limited to fusions that are generally linear, polypeptidic, and located on the termini of Cas9. A conjugation platform that allows the creation of fusions that are non-polypeptidic (e.g., nucleic acids, small molecules, polyethylene glycol [PEG] chains), unnatural peptides/proteins, internally located on Cas9, and branched with a multivalent display, would provide a greater diversity of technologies and applications. For example, precise sequence alteration at the Cas9 cleavage site requires the efficient incorporation of exogenously supplied single-stranded oligonucleotide donor DNA (ssODN)^4^ *via* the homology-directed repair (HDR) pathway.^5–7^ However, most cells instead employ the non-homologous end-joining (NHEJ) repair, which results in unpredictable insertions and deletions of bases at the cleavage site, some of which are large enough to have pathogenic consequences.^3,8,9^ Displaying ssODNs on Cas9 can increase their local concentration around the DNA strand break site to allow enhanced incorporation of the desired sequence. In another application, appending PEG chains to Cas9 may reduce the immunogenicity of this bacterial protein.^10^ Multivalent display of DNA repair pathway inhibitors (e.g., NHEJ or p53 pathway inhibitors) or cell-specific ligands could also enhance precision and efficacious genome editing.^11,12^

An ideal conjugation platform for Cas9 should have several characteristics. First, the platform should be compatible with a diverse set of cargoes (e.g., small molecules, nucleic acids, nanoparticles, antibodies, PEG chains) and allow their multivalent display. Second, the platform should be robust and implementable by non-specialists, given the diverse users of CRISPR technologies. Third, since some of the cargoes (e.g., ssODN) are only available in small quantities and are expensive, the conjugation system should work efficiently without requiring large excesses. Ideally, the platform should be modular and inexpensive to allow for screening multiple conditions (e.g., ssODN sequence). Finally, for real-world applications, the platform should allow scaled-up production of the conjugates following good manufacturing practice (GMP) regulations.^13^

Herein, we present the development of such a platform that relies on thiol-maleimide chemistry and DNA-base pairing, which are both simple, well-established, scalable, and amenable to a wide range of substrates.^14^ Following a structure-guided approach, we systematically scanned the domains of Cas9 to choose residues replaceable with engineered cysteines, to which molecules of any size could be efficiently appended without the loss of Cas9 activity. We successfully appended biotin (small) and PEG groups (large) using thiol-maleimide chemistry. Because many possible conjugates (e.g., ssODNs) are prohibitively expensive for or unamenable to this direct thiol-maleimide conjugation, we also sought to develop a more general conjugation platform. Thus, we designed a short oligonucleotide handle ‘adaptor’, which is attached to Cas9 via thiol-maleimide chemistry and uses base pairing to anchor any molecule containing or appended with nucleic acids (Figure 1a). As an example of platform’s utility, we used it to hybridize long ssODNs that are large, expensive, and not available in sufficient quantities for thiol-maleimide conjugation to Cas9.^15^ The resulting Cas9:ssODN conjugates robustly enhanced the precise incorporation of the desired sequence from ssODN in multiple cell types and genomic sites. Importantly, the chemical conjugation platform enabled the multivalent display of ssODNs, which further enhanced the precise incorporation of the desired sequence over that of the univalent display.

**Figure 1.**
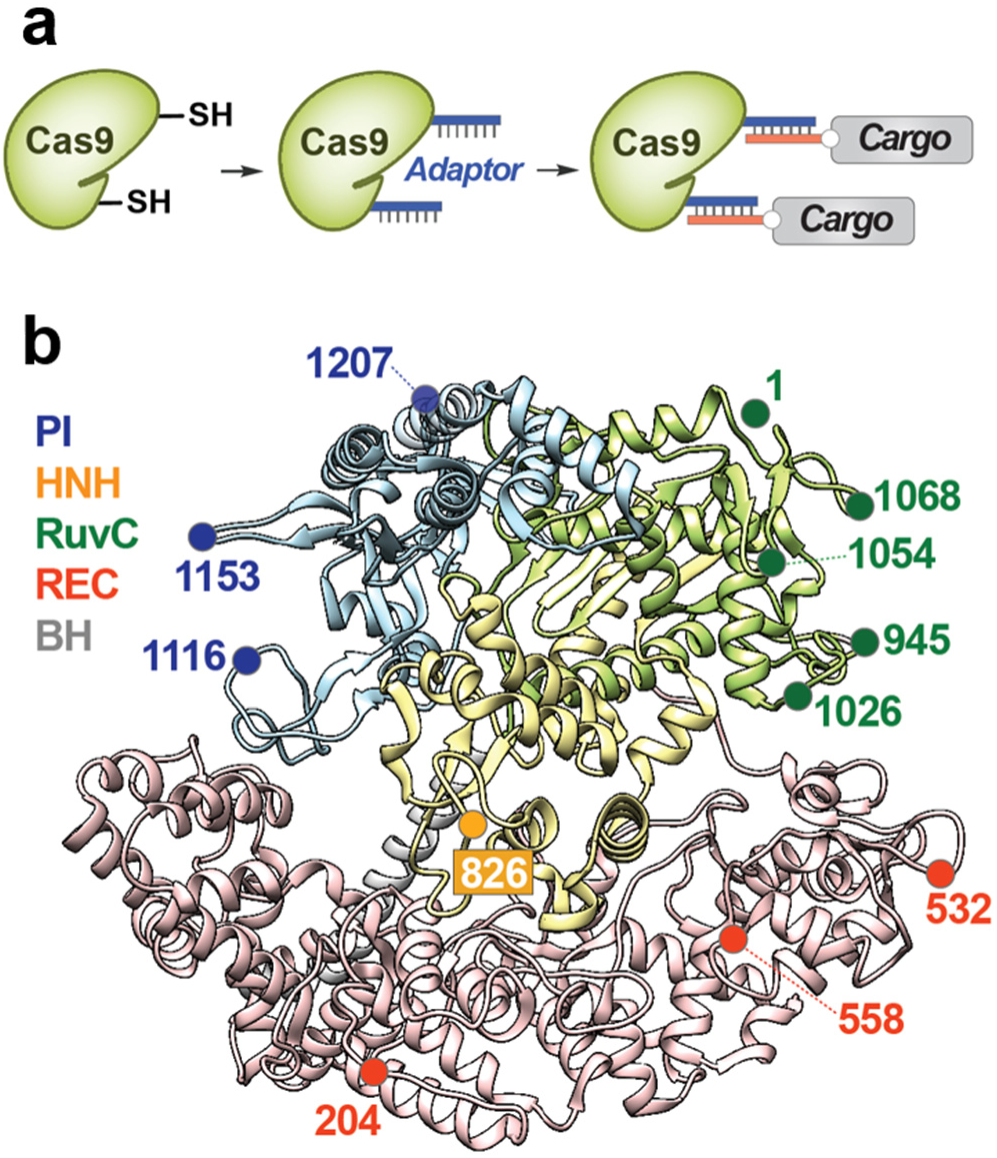
A conjugation platform for Cas9. (a) A modular design strategy to functionalize Cas9. (b) Structure-guided selection of chemical labeling sites. PI domain is in blue, HNH domain is in yellow, RuvC domain is in green, REC domain is in red, and BH is in gray. Crystal structure of the Cas9-gRNA-DNA ternary complex is used (PDB: 5F9R).^28^

Next, we demonstrated the utility of our conjugation platform by efficiently engineering insulin-producing β cells to secrete non-endogenous molecules, including an immunomodulatory protein (~160 residues), without incorporation of any viral or foreign sequences (e.g., promoter) other than that of the secreted molecule. Current β-cell transplantation therapies for type 1 diabetes suffer from immune rejection, resulting in acute cell loss and only short-term therapeutic effects.^16,17^ The macroencapsulation of β cells with a semipermeable membrane can protect them from the host’s immune system, though foreign body reaction-induced fibrosis can impair the mass transfer and viability of encapsulated cells.^18–20^ Anti-inflammatory cytokines, such as interleukin 10 (IL-10), can reduce fibrosis and promotes long-term β-cell survival and superior islet function.^21–25^ Therefore, engineered β-cells that secrete anti-inflammatory cytokines and anti-fibrotic factors will propel the development of cell-based therapeutics for diabetes. By hijacking the insulin expression and secretion machinery, we used our herein developed conjugation platform to efficiently engineer β cells to secrete a non-endogenous peptide in a glucose-responsive manner as is observed for insulin. Furthermore, we were able to engineer β cells to secrete IL-10, demonstrating the immediate usefulness of our conjugation platform for the development of cell-based therapeutics.

## RESULTS

### Multiple domains on Cas9 are tolerant of displaying molecules of diverse size and nature

To choose the sites for conjugation to Cas9, we analyzed the structures of apo-Cas9, gRNA-bound Cas9, and gRNA- and DNA-bound Cas9 for residues that could provide a high labeling yield, tolerate chemical modifications, span all the domains of Cas9, and are surface-exposed in various Cas9 conformations, for efficient display of modifications (Figure 1b).^26–28^ Using aforementioned criteria, we identified two sites (204, 532) on the REC domain, one site (826) on the HNH domain, five sites (1, 945, 1026, 1054, 1068) on the RuvC domain, and two sites (1153, 1207) on the PI domain. We selected residues 558 and 1116 as controls since modifications at 558 will impede the Cas9:gRNA interaction and at 1116 will impede protospacer adjacent motif (PAM) recognition by Cas9 (Figures 1b and S1). We optimized the conjugation conditions for Cas9 variants using biotin-maleimide and PEG (5 kDa)-maleimide as model compounds to ensure that modifications of various sizes or nature were tolerated (Figure S2). The reactions were fast and high yielding at all sites except for the 1153C mutant; sites proximal to 1153C (i.e., 1154C) also yielded low conjugation efficiencies (Figure S2). The location of these residues was not assigned at the crystal structure of apo-Cas9, but the residues were assumed to be amenable to efficient conjugation, since they were expected to be surface-exposed and flexible.^26–28^ Our labeling results, however, indicate that the loop may have higher-order structures that prevent efficient chemical reactions, so we did not use those sites in future experiments. To improve the compatibility of the system with a broader range of conjugates, we next utilized the optimized reaction conditions to label Cas9 at the remaining sites with a 17-nucleotide (nt) DNA adaptor (5’-GCTTCACTCTCATCGTC-3’). The conversion rates were comparable to those of PEG labeling (Figure S2), demonstrating that efficient conjugation of multiple cargo types can occur at these sites. Thus, the identified sites provide high conjugation yields with diverse molecules, including small molecule and polymers (DNA or PEG).

To identify sites that are tolerant to the conjugation of the DNA adaptor without the loss of Cas9 activity, we designed an ssODN that would insert a 33-nt DNA fragment (*HiBiT* sequence^29^) at the target gene (Figure S3). This insertion would result in the expression of a fusion protein with a C-terminal HiBiT tag, which is a small fragment of the NanoLuc luciferase. When HiBiT is complemented by LgBiT, the remainder of NanoLuc, the full-length luciferase is reconstituted to generate a luminescence signal proportional to the degree of knock-in, providing an easy readout for HDR (Figure S3a). We chose *GAPDH* as the first target gene (Figure S3b) owing to its abundant expression in many cell types, which should allow for the reliable detection of the luminescence signal. Using the *HiBiT* knock-in assay, we measured whether appending the DNA adaptor to the cysteine affected Cas9 activity (Figure 2a). As expected, much of Cas9 activity was lost by control modifications at residues 558 and 1116, indicating the reliability of the *HiBiT* knock-in assay. We identified five sites in total from REC domain (532), RuvC domain (1, 945, 1026), and PI domain (1207) whose activity was largely maintained (>85% of wildtype in U2OS), even after labeling with the 17-nt adaptor. Finally, to investigate the off-target profile of the Cas9-adaptor conjugates, we used an *eGFP* disruption assay with matched gRNA and mismatched gRNAs targeting the *eGFP* gene, in U2OS.eGFP.PEST cells.^30,31^ The Cas9-adaptor conjugate retained the target specificity while maintaining the on-target activity (Figure S4).

**Figure 2.**
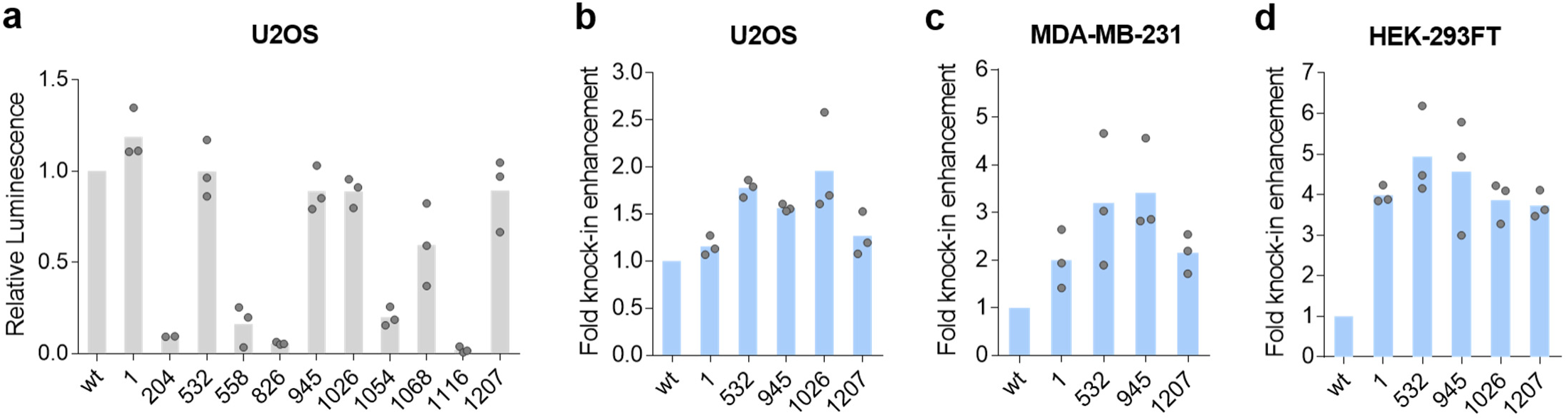
Unitary display of ssODN on Cas9 domains enhances HDR in multiple cell types. (a) *HiBiT* knock-in efficiencies by Cas9-adaptors compared to unlabeled wildtype Cas9 (wt) when a separate Cas9/ssODN system was used. (b-d) ssODN display on Cas9 enhances *HiBiT* knock-in efficiency in various cells: U2OS (b), MDA-MB-231 (c), and HEK-293FT (d). Unlabeled wildtype Cas9 (wt) and Cas9-adaptors labeled at the indicated residues were used. All data from biological replicates are shown.

### Unitary display of ssODN on multiple Cas9 domains enhances HDR in several cell types

Next, we designed ssODN with a sequence complementary to the conjugation adaptor and confirmed the binding of the ssODN bearing the complementary sequence to the Cas9-adaptor using a gel-shift assay (Figure S5). To measure the ability of ssODN conjugates to enhance HDR and site-dependence of such enhancements, we performed the *HiBiT* knock-in assay in U2OS cells. Using the luminescence signals from unconjugated ssODN as normalization controls, we observed enhanced knock-in efficiency at multiple sites (Figures 2b and S6a) when Cas9 displayed ssODN. We were able to confirm such enhancements in multiple cell lines, with a greater than fourfold increase in HEK-293FT cells (Figures 2c, d and S6b, c). For cells with higher HiBIT signal but lower HDR enhancements, we observed site dependence, with two internal conjugation sites (532, 945) generally performing better than the terminal conjugation site (1). An examination of the crystal structure^28^ indicates that cargoes on the two internal residues are expected to align with substrate DNA, while cargoes on the terminal residue project outward from the DNA, which may explain the differences in the HDR-enhancing capacities of different ssODN-bearing sites.

### The ssODN display platform allows rapid and facile screening of multiple conditions

To demonstrate the modular nature of our conjugation platform that should allow rapid testing of multiple conditions and to confirm the generality of HDR enhancement by ssODN display, we tested the ability of the conjugates to enhance HDR under several scenarios (e.g., different genomic sites, ssODN sequences, or readouts). Using the *HiBiT* knock-in assay, we confirmed HDR enhancements at another DNA cleavage site on the *GAPDH* locus (Figures 3a and S7a) and at multiple genomic loci (*PPIB*, *CFL1*; Figures 3a and S7b, c). We then demonstrated HDR enhancement using a fluorescent readout and a longer knock-in fragment (*GFP11*, 57 nt). The correct incorporation of this fragment generated detectable fluorescence through the expression of a fusion protein with a C-terminal GFP11 tag, which forms a fully functional GFP when complemented by GFP1-10 (Figure S8).^32^ Here as well, ssODN display on Cas9 increased the knock-in efficiency by more than threefold (Figures 3b and S7d). In addition to luminescence and fluorescence readouts to demonstrate HDR enhancement, we used a previously reported droplet digital PCR (ddPCR) assay that employs probes to distinguish between wildtype, NHEJ-edited, and HDR-edited sequences at the *RBM20* locus (Figure S9).^33^ All Cas9:ssODN conjugates increased the ratio of HDR over NHEJ, again indicating the generality of our platform (Figure 3c). The conjugates also enhanced HDR when another gRNA/ssODN pair was employed to introduce the same mutation (Figure S10).

**Figure 3.**
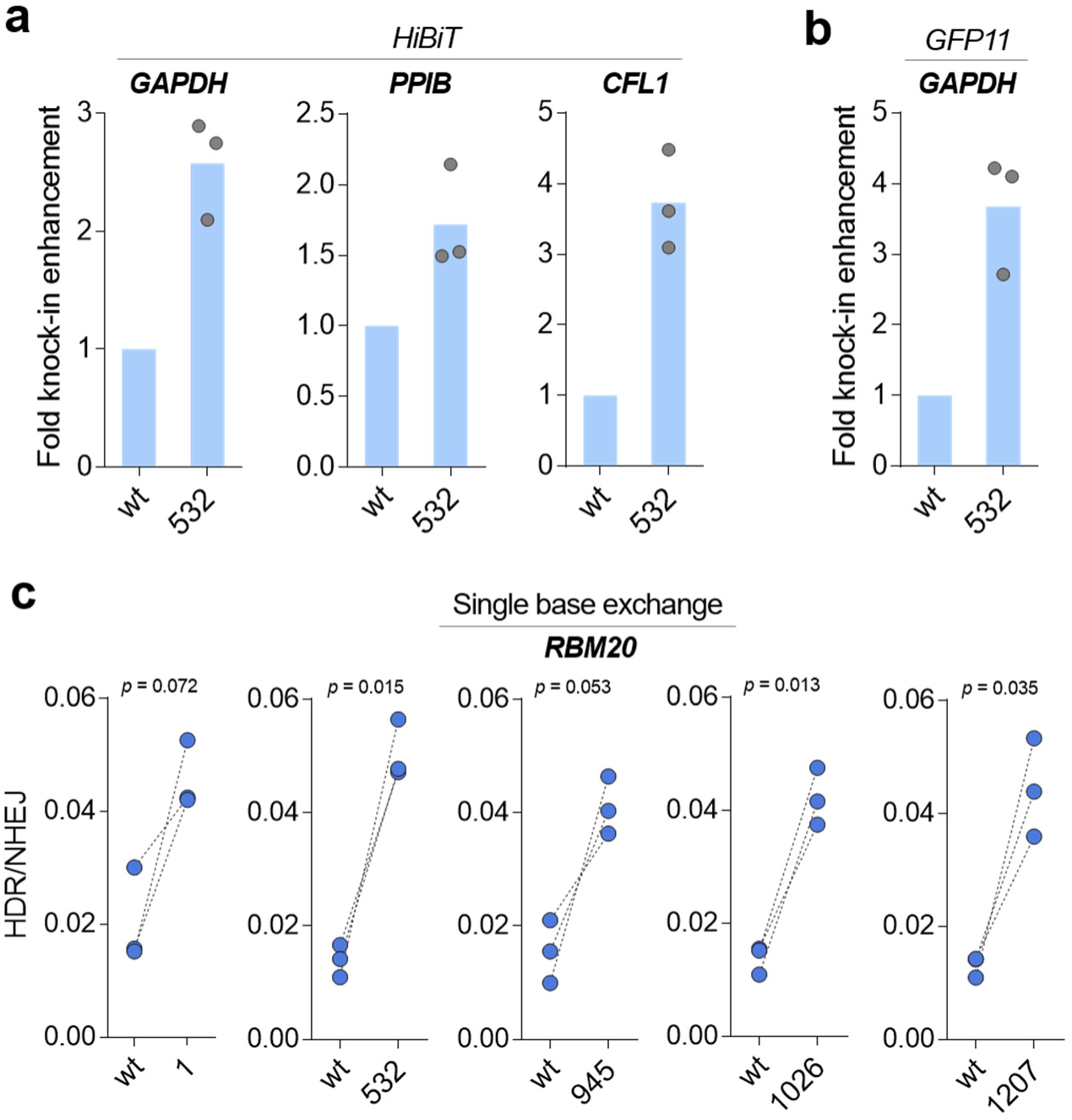
ssODN display platform allows facile testing of multiple conditions. (a) *HiBiT* sequence knock-in efficiency was increased at multiple genomic loci in U2OS or HEK-293FT cells. (b) The *GFP11* sequence insertion at the *GAPDH* locus was promoted in HEK-293T cells. (c) Single base exchange at the *RBM20* locus was promoted in HEK-293FT cells. Unlabeled wildtype Cas9 (wt) and Cas9-adaptors labeled at the indicated residues were used. All data from biological replicates are shown. *P*-values were calculated by paired two-tailed t-test.

### Multivalent display of ssODN on Cas9 further enhances HDR over that of the univalent display

Owing to the small size of our adaptor and the chemical nature of our platform, multivalent displays are feasible (Figure 4a). To demonstrate the multivalent display, we produced Cas9 double-cysteine mutants (532C/945C and 532C/1207C) and labeled them with the adaptor (Figure S11a). Next, we confirmed the binding of the ssODNs to Cas9 (Figure S11b) and observed a boost in HDR efficiency for both the 33-nt *HiBiT* insertion and a single nucleotide exchange (Figures 4b, c, d), indicating that multivalent Cas9 internal modifications further improve the functionality of conjugated Cas9 proteins. To see if an even smaller labeling footprint was possible, we investigated the possibility of further decreasing the length of the adaptor. We found that hybridization by 13 nt or 15 nt showed a similar HDR-enhancing effect as the standard 17-nt pairing (Figure S12), providing greater flexibility to the system.

**Figure 4.**
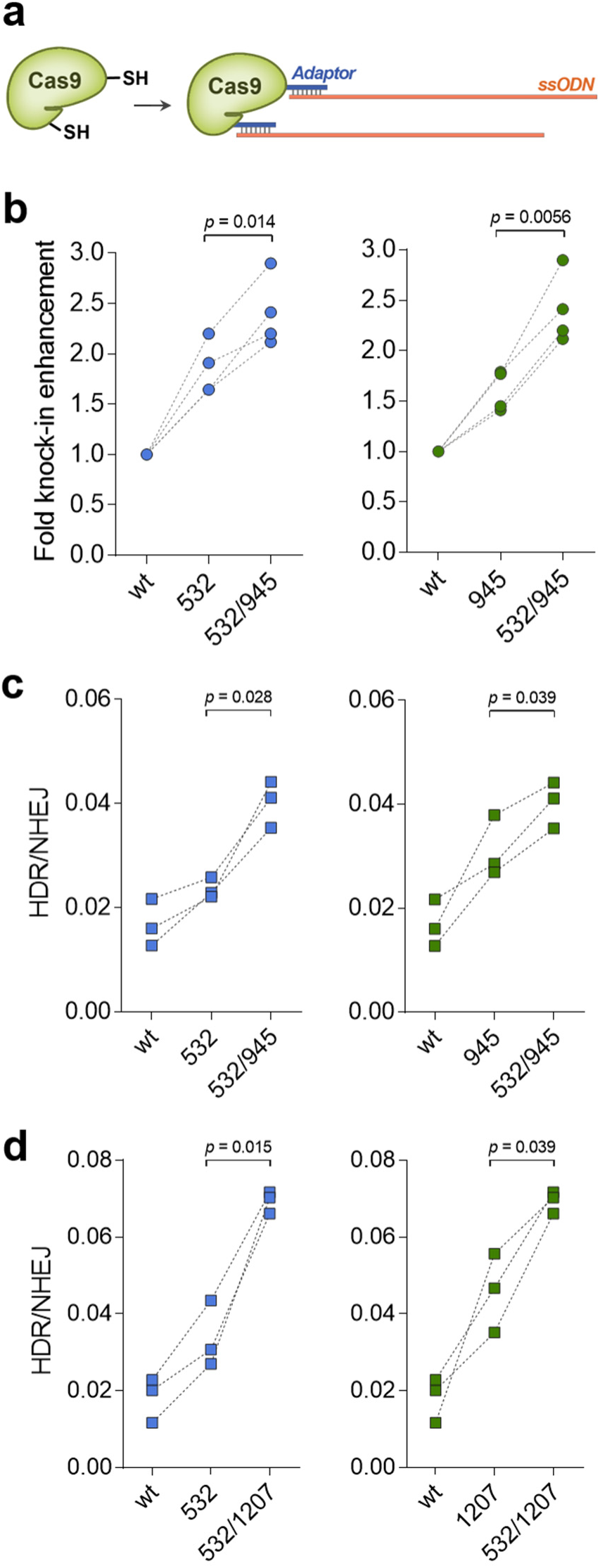
Multivalent display of ssODN further enhances HDR efficiency. (a) Schematic illustrating the production of Cas9 double-ssODN conjugates. (b) *HiBiT* sequence knock-in at the *GAPDH* locus was detected in U2OS cells. (c), (d) Single-nucleotide exchange at the *RBM20* locus was detected in HEK-293FT cells. All data from biological replicates are shown. *P*-values were calculated by paired two-tailed t-test.

### Cas9:ssODN conjugates allow efficient engineering of β cells to secrete IL-10

To demonstrate the functional applicability of our chemically modified Cas9, we used it to efficiently engineer β cells to endow the cells with immunomodulatory function. Since C-peptide is cleaved during proinsulin processing and is co-secreted with insulin, we hypothesized that knocking in the desired gene into the C-peptide portion of the proinsulin locus would enable the secretion of the inserted gene product. Previously, a lentiviral vector encoding a proinsulin-luciferase fusion construct, containing a luciferase inserted into the C-peptide, expressed functional luciferase in levels directly proportion to insulin when stably integrated into the INS-1E β-cell line and responded sensitively to external stimuli, such as glucose concentration.^34^ However, viral-vector engineering poses safety issues such as immunogenicity to viral components or the unintended random insertion of DNA fragments into the host genome.^35,36^ Thus, direct knock-in of the desired gene fragment into the C-peptide using Cas9 will not require regulatory elements (e.g., promoters) and allow co-secretion of the target gene product with insulin without raising immunogenicity issues (Figures 5a and b).

**Figure 5.**
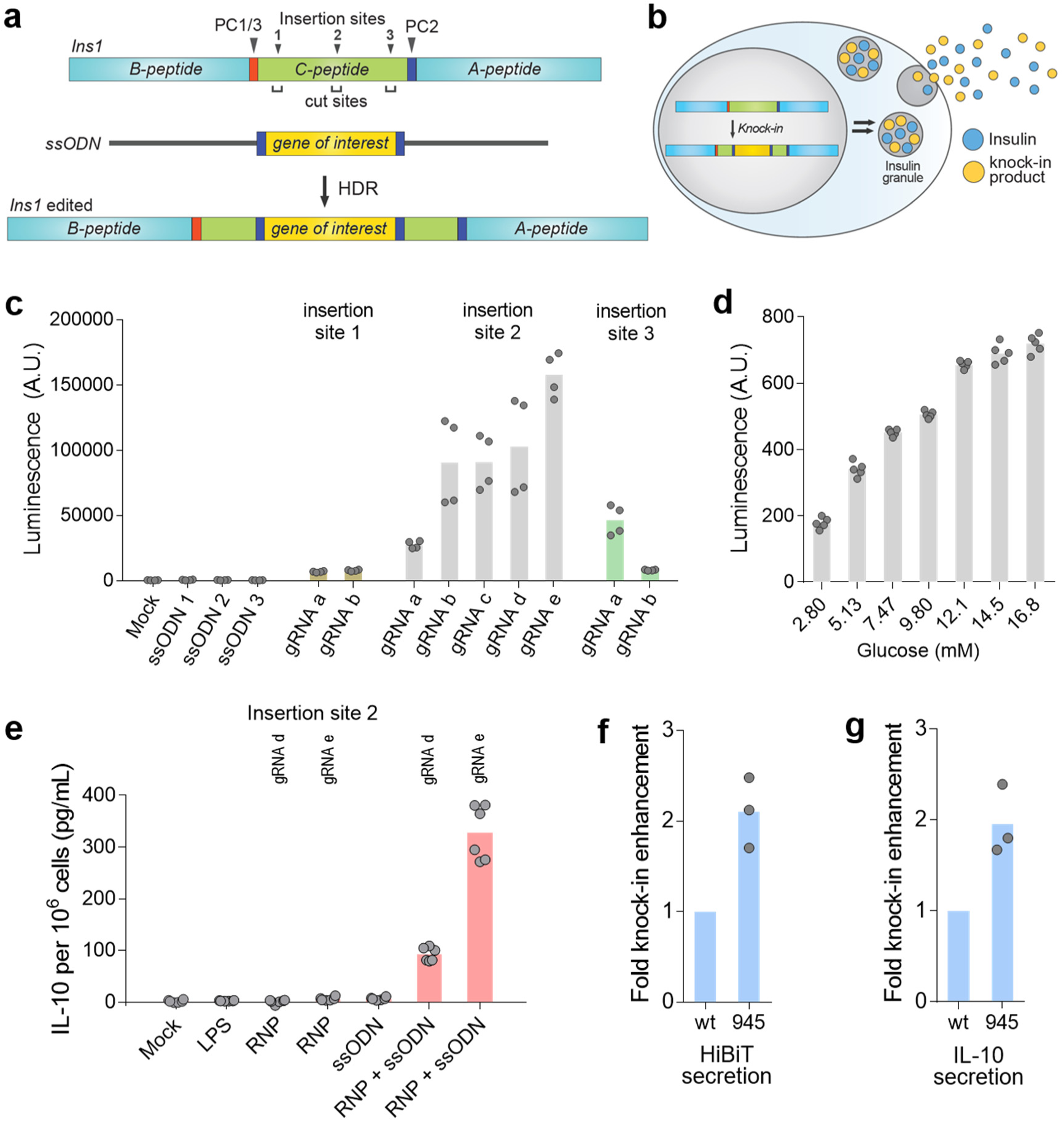
Efficient engineering of INS-1E cells for secretion of exogenous peptides and proteins. (a) Schematic of genome editing in the *Ins1* locus of INS-1E cells. (b) Engineered cells can secret exogenous gene products together with insulin. (c) INS-1E cells were engineered to secrete the 11-residue HiBiT peptide. Multiple gene insertion sites and DNA break sites were investigated. All data from two biological replicates are shown. (d) Glucose-stimulated HiBiT peptide secretion demonstrates knock-in at the *Ins1* locus. All data from five technical replicates are shown. (e) INS-1E cells were engineered to secret IL-10. All data are from two biological replicates are shown. (f-g) Display of ssODN on Cas9 enhanced the secretion of HiBiT peptide (f) and IL-10 (g). All data from biological replicates are shown.

To identify the appropriate insertion site in the *Ins1* locus, we used HDR-mediated knock-in of the *HiBiT* sequence at the C-peptide portion in INS-1E cells (Figure 5a). The target *HiBiT* sequence was flanked by additional prohormone convertase 2 (PC2) cleavage sites^34^ to ensure that no extra amino acids would be present at each end of the knock-in product after processing (Figure 5a). We chose three gene insertion sites at the start, middle, and terminal regions of the C-peptide locus, and designed several gRNAs to target these sites such that insertion sites and DNA cleavage sites would be close enough to obtain high HDR efficiency (Figure 5a). In addition, genome-wide off-target profiles of gRNAs were considered such that potential off-target sites had mismatches at the seed sequences or at least three mismatches in the gene-encoding regions. When standard genome editing was performed at the target sites using non-conjugated Cas9 and ssODN, the HiBiT peptide was secreted from INS-1E cells, which could readily be detected through luminescence signals from the cell culture supernatant after complementation by the LgBiT protein. The highest knock-in efficiency was achieved by targeting the middle region of the C-peptide (site 2) (Figure 5c), so this insertion site was used for future experiments. HiBiT peptide secretion was also stimulated by glucose, indicating that the knock-in product was secreted through the insulin processing and secretion pathways (Figures 5d and S13).

Based on this optimized design, we next knocked-in *Il10*, whose 797-nt ssODN is much larger than that of *HiBIT* (183-nt). The secretory signal peptide sequence present in the *Il10* gene was omitted as our approach leverages the insulin secretion pathway. PC2 cleavage sites were added at each end of *Il10* to obtain intact IL-10 as the knock-in product, and the corresponding ssODN was synthesized by reverse transcription. When INS-1E cells were transfected with both unconjugated Cas9 and ssODN, IL-10 was secreted into the cell culture media as determined *via* enzyme-linked immunosorbent assay (ELISA). No IL-10 was detected after transfection with Cas9 or ssODN alone, or in lipopolysaccharides (LPS)-treated cells^37^ (Figure 5e). We confirmed the correct insertion of the *Il10* gene at the *Ins1* C-peptide region using Sanger sequencing (Figure S14). Finally, we displayed ssODN on Cas9 and found that both HiBiT secretion and IL-10 secretion were significantly promoted by Cas9-ssODN conjugation over that of separate Cas9 and ssODN (Figures 5f, g and S15).

## DISCUSSION

We describe a simple, scalable, and modular chemical platform for site-specific Cas9 labeling with a wide range of functional molecules to expand Cas9 functionality for novel applications. We first identified multiple internal residues on Cas9 that can be modified using thiol-maleimide chemistry without compromising Cas9 activity in cells, opening up a variety of new site-specific conjugation locations with simple and easy-to-use chemistries. We showed that internal conjugation sites and multivalent conjugations often had improved knock-in efficiencies, indicating the need for conjugation systems that are not limited to single modifications of the Cas9 termini. The identified sites could also be used to display inhibitors of DNA repair pathways for their local inhibition at the site of the Cas9-induced double-strand break. For example, co-administration of Cas9 with small-molecule inhibitors of the NHEJ pathway can enhance precision editing,^11^ but concerns about mutagenesis stemming from genome-wide NHEJ inhibition has limited the utility of such inhibitors.^38,39^ Local NHEJ-pathway inhibition at the strand break site through the multivalent display of NHEJ inhibitors on Cas9 itself may allay such concerns. Such local inhibition has been demonstrated for base editors that display peptidic inhibitors of uracil DNA glycosylase for local enzyme inhibition, which improves base-editing efficiencies.^40^ Similarly, the local inhibition of p53 pathway activation can increase the efficiency of precision genome editing in primary cells and stem cells where Cas9-induced double-strand breaks lead to apoptosis *via* this pathway.^41,42^ Finally, displaying tissue-specific ligands on Cas9 will enable cell-specific genome editing.^12^

We also developed a short oligonucleotide handle as a universal anchoring point for any oligonucleotide-containing functional molecules, making this platform amenable to most desired conjugates. When ssODN was attached to this anchor, HDR efficiency was enhanced in multiple cells and genomic loci, simultaneously demonstrating the utility of the conjugation technique as well as the usefulness of an increased local concentration of ssODNs for precision genome editing. Our Cas9-adaptor design is modular in that the Cas9-bearing the universal adaptor can be used for any ssODN or any type of knock-in (e.g., single nucleotide exchange, short DNA insertion, and long gene insertion).

While these studies were underway, reports of genetic fusions of Cas9 to avidin, SNAP tag, or porcine circovirus 2 protein (PCV), which can also bind to donor DNAs, appeared in the literature,^43–46^ and our studies complement these approaches in multiple ways. First, at 17-nt, our Cas9-adaptor constructs are much smaller than the reported constructs, with the possibility of reducing this to as low as 13 nt. Second, while these other genetic fusions are mostly tested at the N- and C-termini of Cas9, we systematically investigated both terminal and internal conjugation sites and found that internal sites yielded higher knock-in efficiencies in certain cell lines. The ability of our system to equally address internal or terminal sites makes it significantly more flexible and adaptable to specific applications. Third, our adaptor-based conjugation strategy does not require chemical modification of the ssODNs, as opposed to avidin- or SNAP-based methods, which can be particularly costly and time-consuming when multiple ssODNs or conditions are being screened during optimization. Fourth, our adaptor sequence can be readily altered to prevent secondary structure formation depending on the ssODN sequence, while the PCV recognition sequence cannot be changed. Finally, owing to the small size of our adaptor and the chemical nature of our platform, multivalent displays are feasible that further enhance HDR and can open up new applications for conjugated Cas9 proteins, expanding the possible range of genome engineering technology.

Novel strategies for β-cell genome editing are urgently needed for endowing the cells with new functions, including immunomodulation, to propel the development of cell-based therapeutics for type 1 diabetes.^47^ Using CRISPR-Cas9 and HDR-based genome editing, we demonstrated that INS-1E cells can be precisely engineered to secret a diverse set of functional molecules, such as small peptides and large gene products. Specifically, we successfully produced cells that co-secrete IL-10, a well-established anti-inflammatory factor that reduces fibrosis and can protect β cells from pro-inflammatory cytokine-induced cell death.^22–25^ This method should also enable the continuous local production of immunomodulatory factors in β cells for preventing β-cell failure, yet due to its local nature, have minimal systematic effects on the host immune system.^48^ Since insulin is secreted in large quantities by β cells, only a small fraction of cells may require editing to have a therapeutic effect. Our approach of hijacking the insulin expression and secretion machinery significantly reduces the size of the exogenous sequences that needs to be knocked-in, allowing efficient engineering of the β cells using Cas9. Moreover, our precise knock-in strategy would be safer than conventional random gene integration methods using viral vectors that are immunogenic and result in unpredictable genomic sequence.^35,36^ Using this strategy, β cells could also be engineered to secrete glucagon-like peptide 1 (GLP-1), which when co-secreted with insulin may enhance the function and viability of β cells.^49^

Overall, this study provides a simple and effective method for forming chemical conjugates of Cas9 to enhance its functionality, and its ease-of-use will make it convenient for scientists of all backgrounds to modify existing Cas9 tools to suit their desired applications. The example application of our method shows that Cas9:ssODN conjugates can successfully enhance precision genome editing in β cells, opening up a new possibility of chemically enhanced Cas9 in regenerative medicine.

## ACKNOWLEDGEMENTS

This work was supported by the Burroughs Wellcome Fund (Career Award at the Scientific Interface), DARPA (BrdiN66001-17-2-4055), and NIH (UC4DK116255, R01 DK113597, HL095722).

## AUTHOR CONTRIBUTIONS

D.L., K.J.C., and A.C. designed the conjugation platform. D.L. (with assistance from B.K.L) developed the conjugation platform. D.L., V.S., and A.C. designed β-cell engineering experiments with inputs from B.K.W. and J.M.K., and D.L. and V.S. performed those experiments. D.L. and A.C. wrote the manuscript, which was edited by all the authors.

## ADDITIONAL INFORMATION

The supplementary information is available online.

## METHODS

### Cas9 expression and purification

A plasmid for SpCas9 expression (2x NLS and C-terminal His tag, pET-28a) was a gift from the Gao group (Addgene #98158).^50^ *E. coli* Rosette2 (DE3) expressing wildtype Cas9, single-cysteine Cas9 mutants, or double-cysteine Cas9 mutants were grown overnight at 18°C with 0.5 mM of IPTG supplemented when the OD_600 nm_ reached 0.8–1.2. The protein was purified by successive Ni-NTA affinity chromatography, cation exchange chromatography, and size-exclusion chromatography. Purified proteins were snap-frozen in liquid nitrogen and stored at −80°C in Cas9 storage buffer (20 mM Tris·HCl, 0.1 M KCl, 1 mM TCEP, 20% glycerol, pH 7.5).

### Site-directed mutagenesis

Two cysteine residues in SpCas9 (C80, C574) were replaced by serine to give a cysteine-free mutant. Based on this construct, multiple single-cysteine and double-cysteine mutants were generated by introducing cysteines at the designated residues. Mutagenesis was performed using the partial overlapping primer design method or using a Q5 Site-Directed Mutagenesis Kit (New England Biolabs).

### Cas9 labeling by thiol-maleimide conjugation

Adaptor oligonucleotide (GCT TCA CTC TCA TCG TC) modified with protected maleimide (maleimide-2,5-dimethylfuran cycloadduct) at the 5’ terminus was synthesized by Gene Link. Prior to thiol-maleimide conjugation, the maleimide group was deprotected via retro-Diels-Alder reaction by heating the DNA in toluene for 3 h at 90°C. Solvent was removed under the reduced pressure, and the resulting DNA in solid form was dissolved in water to give 2 mM solution. Cas9 cysteine mutants (4 μM) were mixed with 300 μM of PEG (5 kDa)-maleimide or adaptor oligonucleotide-maleimide in reaction buffer (20 mM Tris·HCl, 0.1 M KCl, 1 mM TCEP, pH 7.5). The reaction proceeded for 3 h at room temperature (RT) with mild rotation. The resulting mixture was diluted with a high-salt buffer (20 mM Tris·HCl, 1 M KCl, 1 mM TCEP, 20% glycerol, pH 7.5) and incubated with Ni-NTA agarose beads at 4°C. The beads were extensively washed with the high-salt buffer to completely remove non-specifically bound oligonucleotide molecules. Labeled Cas9 was eluted with an elution buffer (20 mM Tris·HCl, 0.1 M KCl, 1 mM TCEP, 250 mM imidazole, 10% glycerol, pH 7.5). Finally, buffer exchange was conducted using Amicon Ultra-0.5 mL centrifugal filters with a 100 kDa cut-off (Millipore) to give Cas9-adaptor conjugates in storage buffer (20 mM Tris·HCl, 0.1 M KCl, 1 mM TCEP, 10% glycerol, pH 7.5).

### Cas9 biotin labeling and pull-down by streptavidin beads

Cas9 with enhanced specificity [eSpCas9(1.1)]^51^ was used for biotin labeling. Cas9 cysteine mutants (7 μM) were mixed with 500 μM of EZ-Link™ Maleimide-PEG_2_-Biotin (Thermo) in a reaction buffer (20 mM Tris·HCl, 0.1 M KCl, 1 mM TCEP, pH 7.5). The reaction proceeded for 3 h at room temperature (RT) with mild rotation. Excess compounds were removed by Bio-Gel P-6 columns (Biorad) according to the manufacturer’s protocol. Next, 30 pmol of Cas9 from the above step was incubated with 30 μL of Pierce Streptavidin Magnetic Beads (Thermo) overnight at 4°C. Flow-through was collected and the beads were washed twice with a washing buffer (20 mM Tris·HCl, 0.15 M NaCl, 0.1% Tween20, pH 7.4; 300 μL) and once with the reaction buffer (300 μL). The beads were heated to 95°C for 5 min in the presence of SDS-PAGE buffer, and the resulting bead-bound fraction (eluate) and flow-through were subjected to SDS-PAGE followed by Coomassie staining.

### Electrophoretic mobility shift assay

For this assay, 300 nM of Cas9 was mixed with 300 nM of ssODNs in a binding buffer (20 mM Tris·HCl, 0.1 M KCl, 1 mM TCEP, 10% glycerol, pH 7.5). For the Cas9 double-adaptor conjugates, 200 nM of protein and 400 nM of ssODN were used. For testing long ssODNs (Figure S15e), 80 nM Cas9 and 60 nM ssODN were used. The mixture was incubated for 30 min at RT and resolved by 1% agarose gel. DNA was stained using SYBR Gold, and fluorescence images were obtained using an Azure c600 (Azure Biosystems) with the Cy3 channel.

### *In vitro* transcription to synthesize single-guide RNAs

Sequences of target-specific forward primers and universal reverse primers are listed in Table S2. Polymerase chain reactions (PCRs) were conducted using Q5 High-Fidelity 2x Master Mix (New England Biolabs) in the presence of 0.5 µM of forward and reverse primers in a volume of 25 µL. The PCR program was as follows: Initial denaturation at 95°C for 1 min; 25 cycles of 95°C for 15 s, 58°C for 30 s, and 72°C for 15 s; final extension at 72°C for 2 min and cooling to 25°C using a 1% ramp. The resulting mixture was used for *in vitro* transcription without purification. The reaction was performed using the HiScribe T7 Quick High Yield RNA Synthesis Kit (New England Biolabs). The mixture contained 10 µL of NTP buffer mix, 2 µL of the above crude PCR product, 2 µL of T7 RNA polymerase mix, and 0.75 µL (30 U) of recombinant RNase inhibitor (New England Biolabs) in a final volume of 30 µL. The reaction was conducted for 12 h at 37°C. DNase treatment was performed to remove template DNA according to the manual. RNAs were purified using the MEGAclear Transcription Clean-Up Kit (Invitrogen) according to the manual.

### Short single-stranded oligonucleotides

Single-stranded donor DNAs for *HiBiT* insertion, *GFP11* insertion, and single nucleotide exchange at the *RBM20* locus were Ultramer DNA oligonucleotides synthesized by Integrated DNA Technology. Their sequences are listed in Table S3.

### Long single-stranded oligonucleotides for *Il10* insertion

Single-stranded donor DNAs for *Il10* insertion were synthesized by reverse transcription.^52^ First, double-stranded gBlocks DNAs were synthesized by Integrated DNA Technology. The DNAs have the T7 promoter sequences followed by the reverse complementary sequences of the final ssODN sequences. DNAs were produced in large quantities by PCR, followed by gel electrophoresis and gel extraction. Next, *in vitro* transcription was performed using the HiScribe T7 Quick High Yield RNA Synthesis Kit (New England Biolabs). The mixture contained 10 µL of NTP buffer mix, 400 ng of the double-stranded DNA template, 2 µL of T7 RNA polymerase mix, and 0.4 µL (16 U) of recombinant RNase inhibitor (New England Biolabs) in a final volume of 20 µL. The reaction was conducted for 4 h at 37°C. DNase treatment was performed to remove template DNA according to the manual. The resulting RNAs were purified using the MEGAclear Transcription Clean-Up Kit (Invitrogen) according to the manual. Finally, reverse transcription was performed to obtain single-stranded donor DNAs. Approximately 200−250 pmol of RNA was mixed with 400 pmol of reverse primer and 6 μL of dNTP mix (25 mM each, New England Biolabs) in nuclease-free water at a final volume of 35 μL. The mixture was incubated at 65°C for 5 min, then immediately placed on ice for 5 min to induce RNA−primer annealing. Then, 10 μL of 5x RT buffer (250 mM Tris·HCl, 375 mM KCl, 15 mM MgCl_2_, pH 8.3), 2.5 μL of 0.1 M dithiothreitol solution, 0.5 μL (20 U) of RNase inhibitor (New England Biolabs), and 2.5 μL of TGIRT-III reverse transcriptase (InGex) were added to the RNA−primer solution. The reaction was proceed at 58°C for 3 h. Next, RNA was hydrolyzed by adding 21 μL of 0.5 M NaOH solution and heating at 95°C for 10 min. The basic solution was quenched by the addition of 21 μL of 0.5 M HCl solution. The resulting single-stranded DNAs were purified using MinElute PCR Purification Kit (Qiagen) according to the manual. The purity of the ssDNA was confirmed by 6% TBE-Urea gel electrophoresis followed by SYBR Gold staining. All DNA sequences are listed in Table S4.

### *HiBiT* sequence knock-in by nucleofection

U2OS cells stably expressing *eGFP.PEST* or MDA-MB-231 cells were transfected with Cas9 ribonucleoprotein (RNP) and ssODN using the SE Cell Line 4D-Nucleofector kit (Lonza) following the pulse program of DN-100 (U2OS) or CH-125 (MDA-MB-231). For Cas9:ssODN conjugates, 10 pmol of Cas9-adaptor were pre-mixed with 10 pmol of ssODN and incubated at RT for 15–30 min prior to RNP formation to ensure Cas9:ssODN conjugate formation. Then 10 pmol of gRNA was added, and the final mixture was incubated for 5–10 min at RT. In cases where Cas9 did not specifically bind ssODNs, the RNP was formed first because it is known that nonspecific Cas9-DNA interactions hamper the RNP formation. After incubating Cas9 and gRNA at RT for 5–10 min, 10 pmol of ssODN were added to the mixture. Approximately 200,000 cells were transfected with the above mixtures in a well of the nucleofection kit, and 20,000 transfected cells were seeded in each well of a 96-well plate. Cells were incubated for 24 h at 37°C, and cell viability was measured using the PrestoBlue Cell Viability Reagent (Thermo). Next, the HiBiT detection was performed using the Nano-Glo HiBiT Lytic Detection System (Promega) according to the manufacturer’s protocol. The resulting luminescence signals were normalized based on the cell viability.

### *HiBiT* sequence knock-in by lipofection

HEK-293FT cells were seeded in a 96-well plate at a density of 10,000 cells per well. The next day, Lipofectamine CRISPRMAX (Invitrogen) was used to transfect the cells with Cas9 RNP and ssODN, with final concentrations of 25 nM of both reagents in 110 µL of medium per well in a 96-well plate. For Cas9:ssODN conjugates, the Cas9-adaptor was pre-mixed with the ssODN in Opti-MEM (Gibco) and incubated at RT for 15–30 min prior to RNP formation. Next, gRNA was added, and the mixture was incubated for 5–10 min at RT. Then, Plus reagent (Thermo; 0.17 µL per well) was added, and the mixture was incubated for an additional 5 min. Finally, Lipofectamine CRISPRMAX (0.3 µL per well) in Opti-MEM was added, and the mixture was incubated at RT for 5 min. The final transfection mixture was transferred to each well. In cases where Cas9 did not specifically bind ssODNs, the RNP was first formed by incubating Cas9 and gRNA at RT for 5–10 min in Opti-MEM. Then, Plus reagent (0.17 µL per well) was added, and the mixture was incubated for an additional 5 min. Next, the ssODN was added to the mixture. Finally, Lipofectamine CRISPRMAX (0.3 µL per well) in Opti-MEM was added, and the mixture was incubated at RT for 5 min. The final transfection mixture was transferred to each well. The transfections were performed in three technical replicates for each biological replicate. For knock-in experiments using Cas9 double-ssODN conjugates (Figure 5), RNP formation was performed first to prevent the non-specific Cas9-ssODN interaction that blocks RNP formation and decreases the genome editing efficiency. A luminescence detection assay was performed as described above at 24 h post transfection.

### *GFP11* sequence knock-in by lipofection

HEK-293T cells were seeded in a 96-well plate at a density of 8,000 cells per well. The next day, Lipofectamine CRISPRMAX was used to transfect the cells with Cas9 RNP (30 nM) and ssODN (30 nM) using the same procedures as described above for the HiBiT knock-in assay. Approximately 20–22 hours post transfection, the media was exchanged, and the cells were incubated for an additional 2–4 hours. Then, a plasmid encoding for a *GFP1-10* fragment (Addgene #70219, a kind gift from Prof. Bo Huang)^32^ was delivered to the cells using Lipofectamine 2000 (Invitrogen) (120 ng plasmid and 0.4 µL Lipofectamine per well). A total of 50 h after RNP and ssODN transfection, the cells were fixed using 4% paraformaldehyde. Nuclei were stained by HCS NuclearMask Blue Stain (Invitrogen), and fluorescence images were obtained using the ImageXpress Micro (Molecular Devices) with the DAPI and GFP channels.

### Droplet digital PCR-based assay to quantify NHEJ and HDR

HEK-293FT cells were seeded in a 96-well plate at a density of 10,000 cells per well. The next day, the cells were transfected with Cas9 RNP (35 nM) and ssODN (35 nM) using Lipofectamine RNAiMAX (Invitrogen) in 110 µL of media per well in a 96-well plate. For Cas9:ssODN conjugates, the Cas9-adaptor was pre-mixed with ssODN in Opti-MEM (Gibco) and incubated at RT for 15–30 min prior to RNP formation. Next, gRNA was added, and the mixture was incubated for 5–10 min at RT. Finally, Lipofectamine RNAiMAX (0.3 µL per well) in Opti-MEM was added, and the mixture was incubated at RT for 5 min. The final transfection mixture was transferred to each well. In cases where Cas9 does not specifically bind to the ssODNs, the RNP was first formed by incubating Cas9 and gRNA at RT for 5–10 min in Opti-MEM. Then, the ssODN was added to the mixture. Finally, Lipofectamine RNAiMAX (0.3 µL per well) in Opti-MEM was added, and the mixture was incubated at RT for 5 min. The final transfection mixture was transferred to each well. Two days post transfection, the cells were harvested, and the genomic DNA was extracted using a DNeasy Blood & Tissue Kit (Qiagen). Genomic sequences were read by droplet digital PCR as previously reported.^33^ For the single-nucleotide exchange experiments using Cas9 double-ssODN conjugates (Figure 4), RNP formation was performed first because more ssODN was used in comparison to RNP.

### *eGFP* disruption assay to confirm the target specificity of the Cas9-adaptor conjugate

Cas9 (10 pmol) and gRNA (10 pmol) were mixed and incubated at RT for 5 min. U2OS.eGFP.PEST cells were transfected with the RNP complex using the SE Cell Line 4D-Nucleofector kit (Lonza) following the pulse program of DN-130. After transfection, cells were suspended in the culture media and transferred to a 96-well plate (20,000 cell/well). Forty-eight hours after transfection, cells were fixed with 4% paraformaldehyde solution and nuclei were stained with HCS NuclearMask Blue Stain (Invitrogen). The resulting fluorescence signals from eGFP and nuclei were measured using an ImageXpress Micro High Content Analysis System (Molecular Devices).

### *HiBiT* sequence knock-in by nucleofection in INS-1E cells

INS-1E cells were transfected with Cas9 RNP and ssODN using the SF Cell Line 4D-Nucleofector kit (Lonza) following the pulse program of DE-130. For Cas9:ssODN conjugates, 20 pmol of Cas9-adaptor were pre-mixed with 20 pmol of ssODN and incubated at RT for 15–30 min prior to RNP formation to ensure Cas9:ssODN conjugate formation. Then 20 pmol of gRNA was added, and the final mixture was incubated for 5–10 min at RT. In cases where Cas9 did not specifically bind ssODNs, the RNP was formed first because nonspecific Cas9-DNA interactions can hamper the RNP formation. After incubating Cas9 and gRNA at RT for 5–10 min, 20 pmol of ssODN were added to the mixture. Approximately 200,000 cells were transfected with the above mixtures in a well of the nucleofection kit, and cells were seeded in a well of a 24-well plate. Cells were incubated at 37°C for 48 h, and the supernatant was taken to measure the amount of secreted HiBiT peptide using the Nano-Glo HiBiT Extracellular Detection System (Promega). The resulting luminescence signals were normalized based on the cell viability that was measured using the PrestoBlue Cell Viability Reagent (Thermo).

### *Il10* knock-in by nucleofection in INS-1E cells

INS-1E cells were transfected with Cas9 RNP and ssODN as described above for *HiBiT* knock-in. For Cas9:ssODN conjugates, 20 pmol of Cas9-adaptor were pre-mixed with 12 pmol of ssODN and incubated at RT for 15–30 min prior to RNP formation to ensure Cas9:ssODN conjugate formation. Then 20 pmol of gRNA was added, and the final mixture was incubated for 5–10 min at RT. In cases where Cas9 did not specifically bind ssODNs, the RNP was formed first because nonspecific Cas9-DNA interactions can hamper the RNP formation. After incubating Cas9 and gRNA at RT for 5–10 min, 12 pmol of ssODN were added to the mixture. Approximately 200,000 cells were transfected with the above mixtures in a well of the nucleofection kit, and cells were seeded in a well of a 24-well plate. Cells were incubated at 37°C for 72 h, and the supernatant was taken to measure the amount of secreted IL-10 using the IL-10 Rat ELISA Kit (Invitrogen, catalog # BMS629). The resulting values were normalized based on the cell viability. LPS was used at the concentration of 10 µg/mL.

### Glucose-stimulated peptide secretion

INS-1E cells knocked in with the *HiBiT* sequence were grown in a large scale. Then, cells were seeded in a 24-well plate at the density of 150,000 cells per well. The next day, cell were washed with and incubated in Krebs-Ringer bicarbonate buffer (138 mM NaCl, 5.4 mM KCl, 5 mM NaHCO_3_, 2.6 mM MgCl_2_, 2.6 mM CaCl_2_, 10 mM HEPES, pH 7.4, 0.5% BSA) without glucose for 2 h. Cells were subsequently incubated with Krebs-Ringer bicarbonate buffer containing glucose (from 2.8 mM to 16.8 mM) for 1 h. The supernatant was taken to measure the amount of secreted HiBiT peptide using the Nano-Glo HiBiT Extracellular Detection System (Promega).

### PCR to amplify the *Il10* knock-in sequence

Genomic DNAs from the edited INS-1E cells were extracted using a DNeasy Blood & Tissue Kit (Qiagen). PCR was performed using 50 ng of genomic DNA, 0.5 µM of forward primer (CCC GGA GAA GCG TAG CAA AG), 0.5 µM of reverse primer (AAA GAT TCC CGT TCA CAC AAT CC), and Q5 Hot Start High-Fidelity 2x Master Mix (New England Biolabs) in a final volume of 25 µL.

## Supporting Information

### 1. Supplementary tables and figures

**Table S1.**
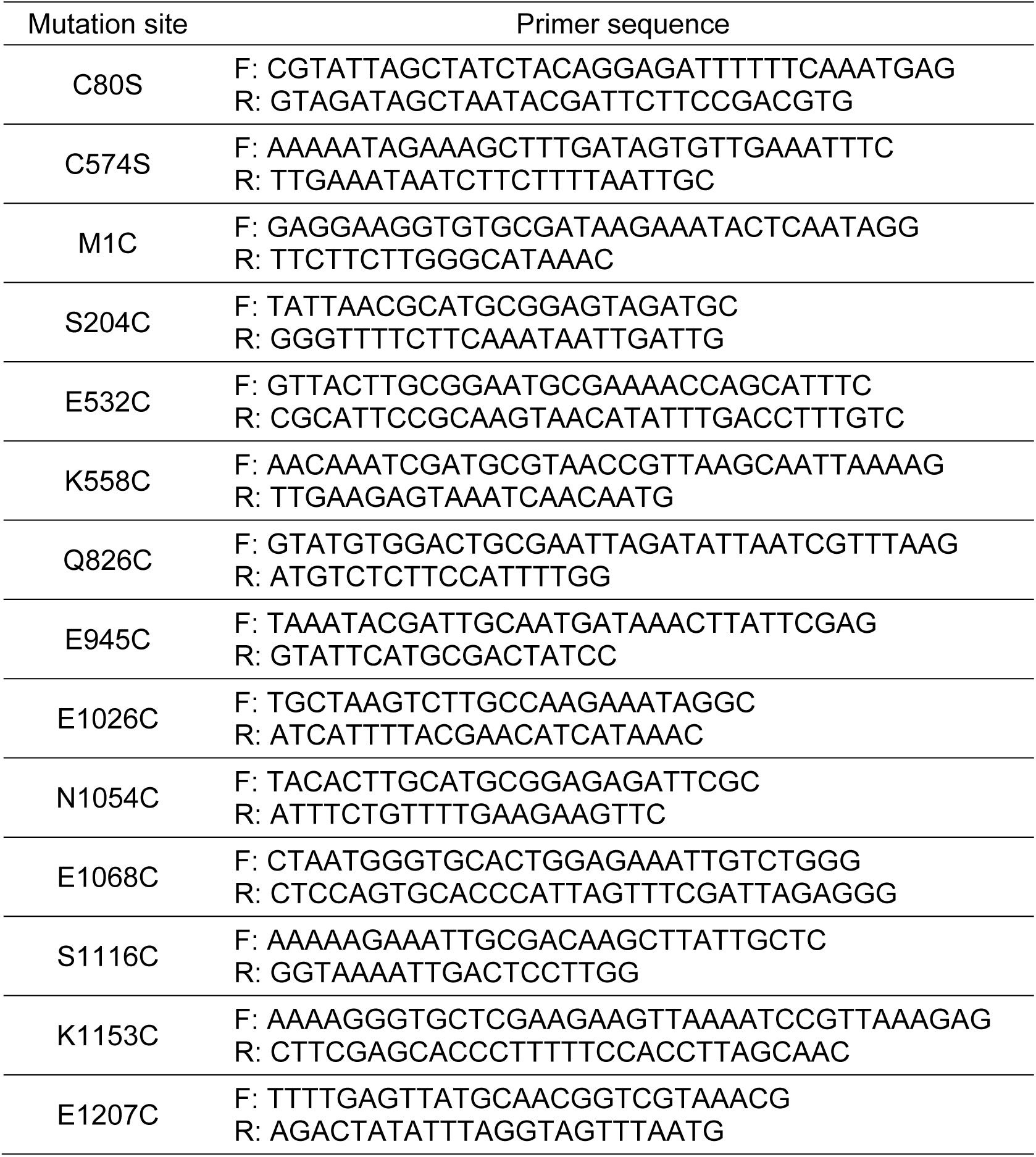
Primer sequences for mutagenesis.

**Table S2.**
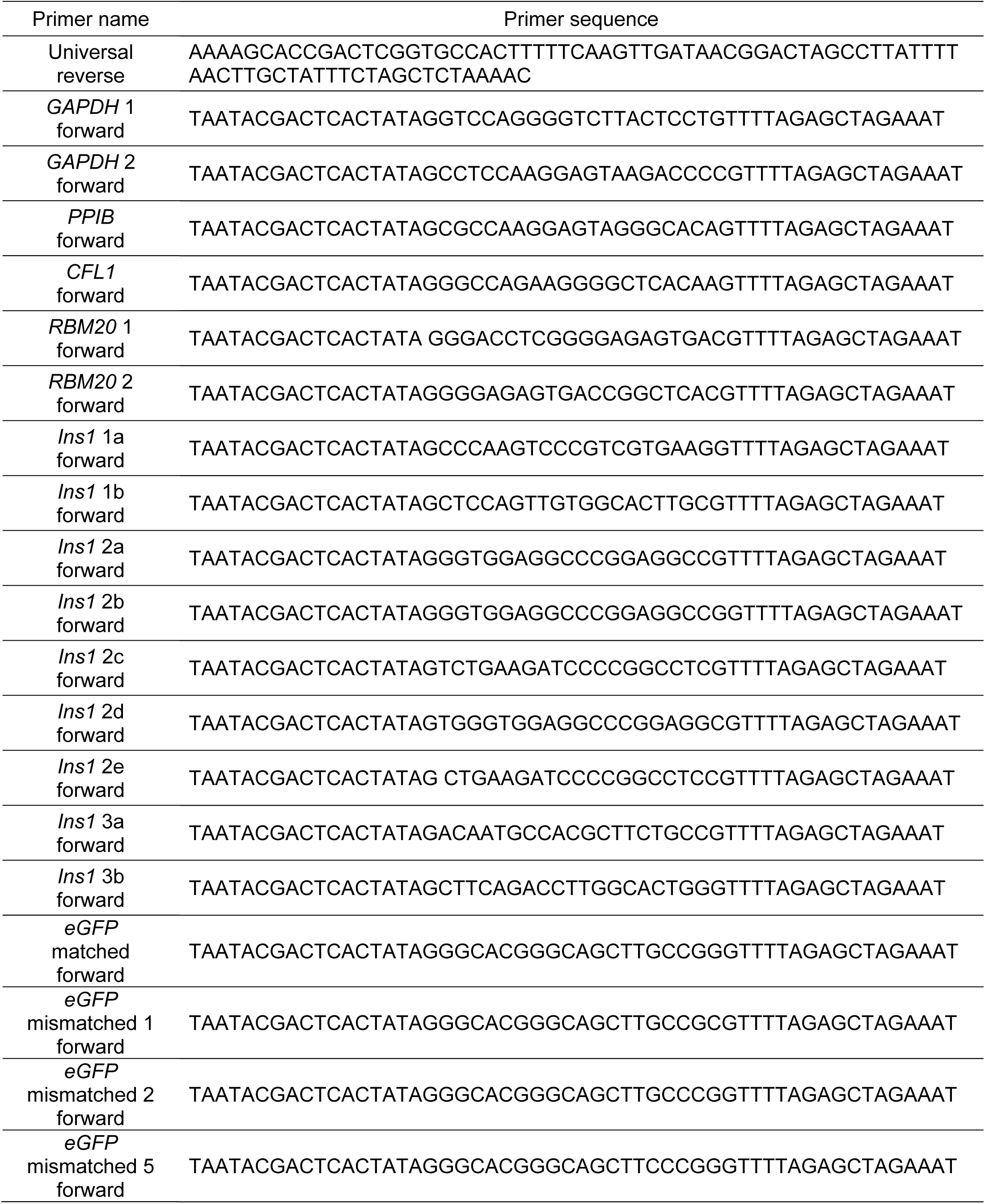
Primer sequences for gRNA synthesis.

**Table S3.**
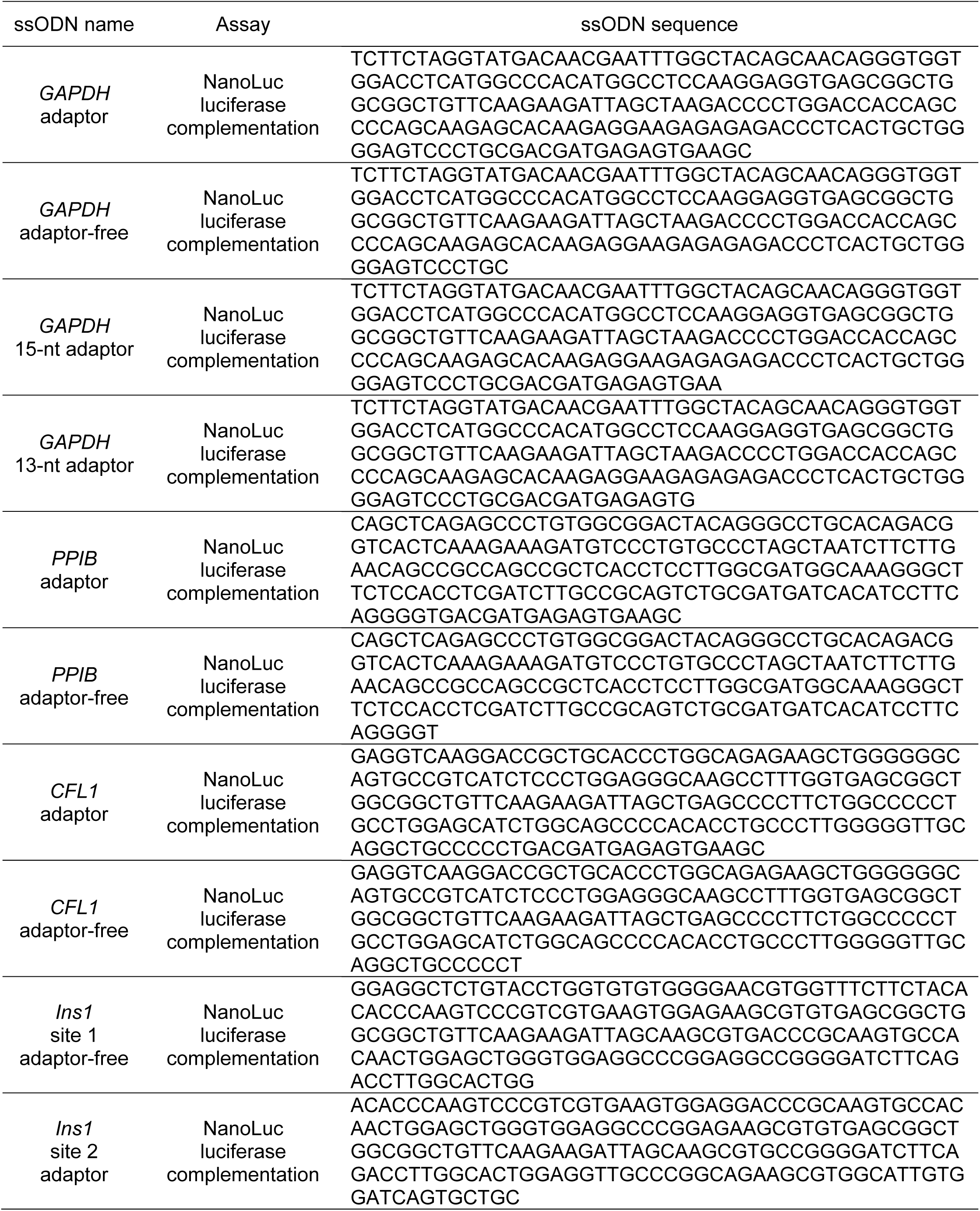
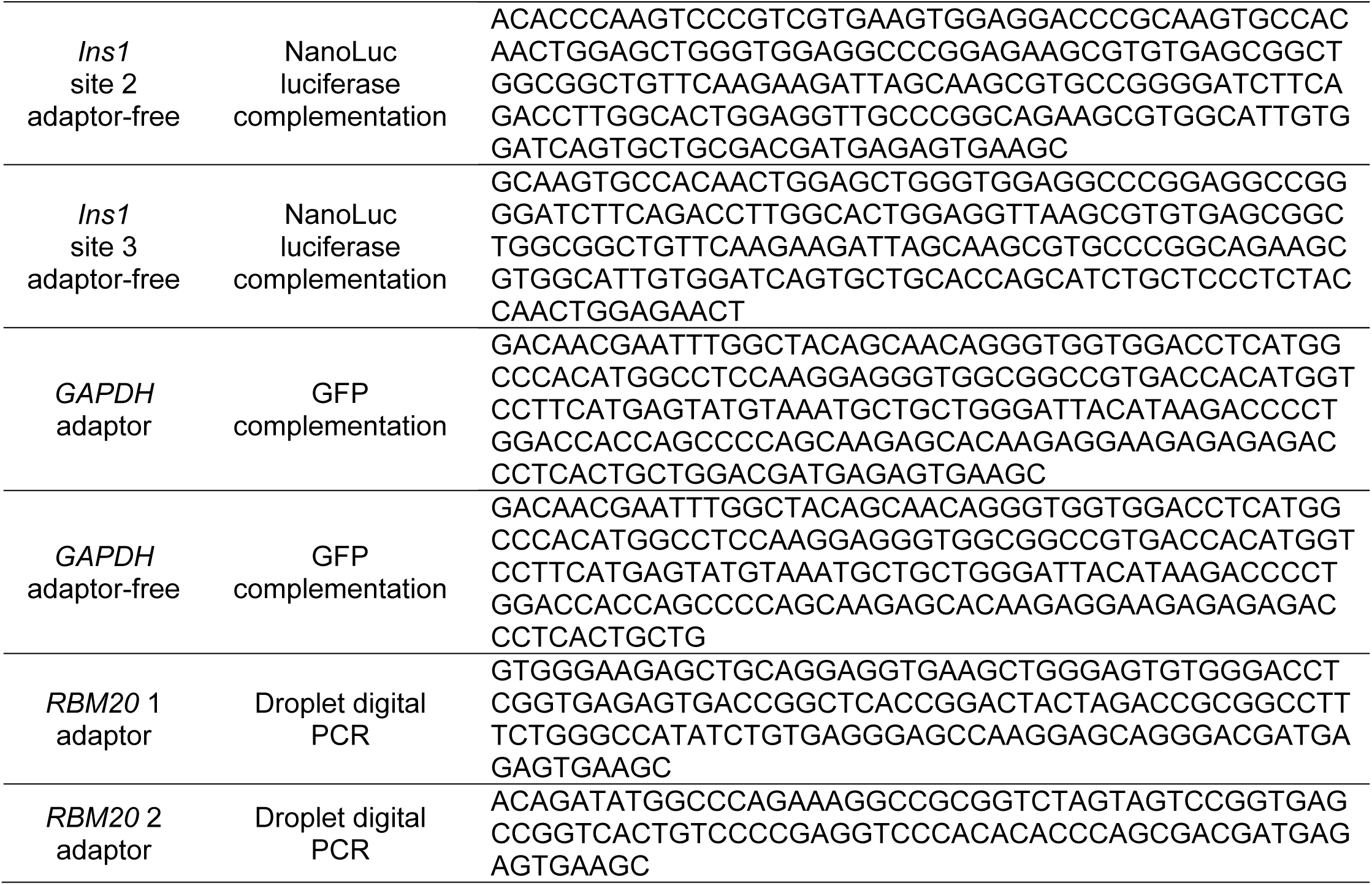
Sequences of ssODNs (< 200 nt).

**Table S4.**
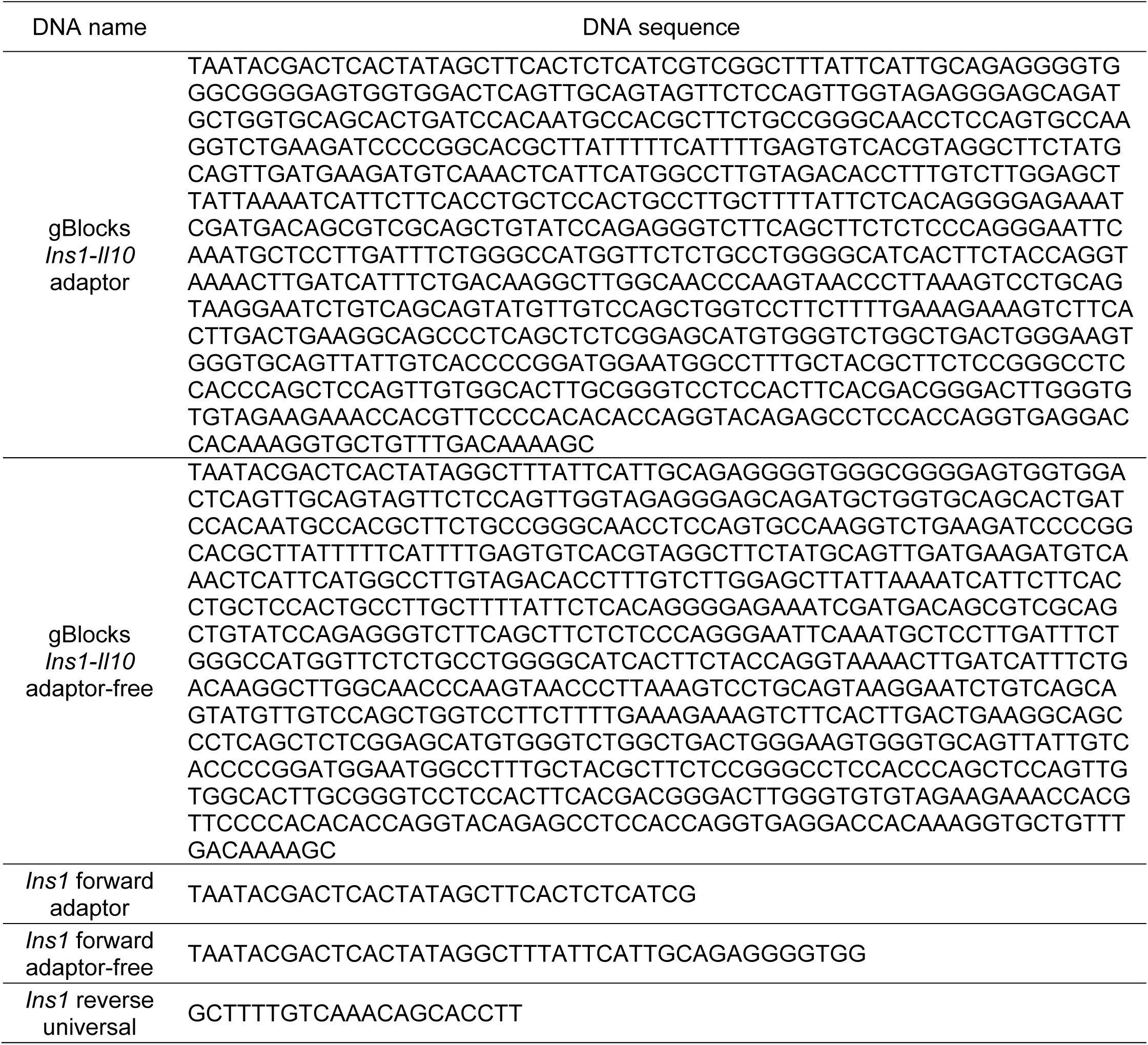
Sequences of gBlocks DNAs and primers for generating ssODNs for *IL-10* knock-in.

**Table S5.**
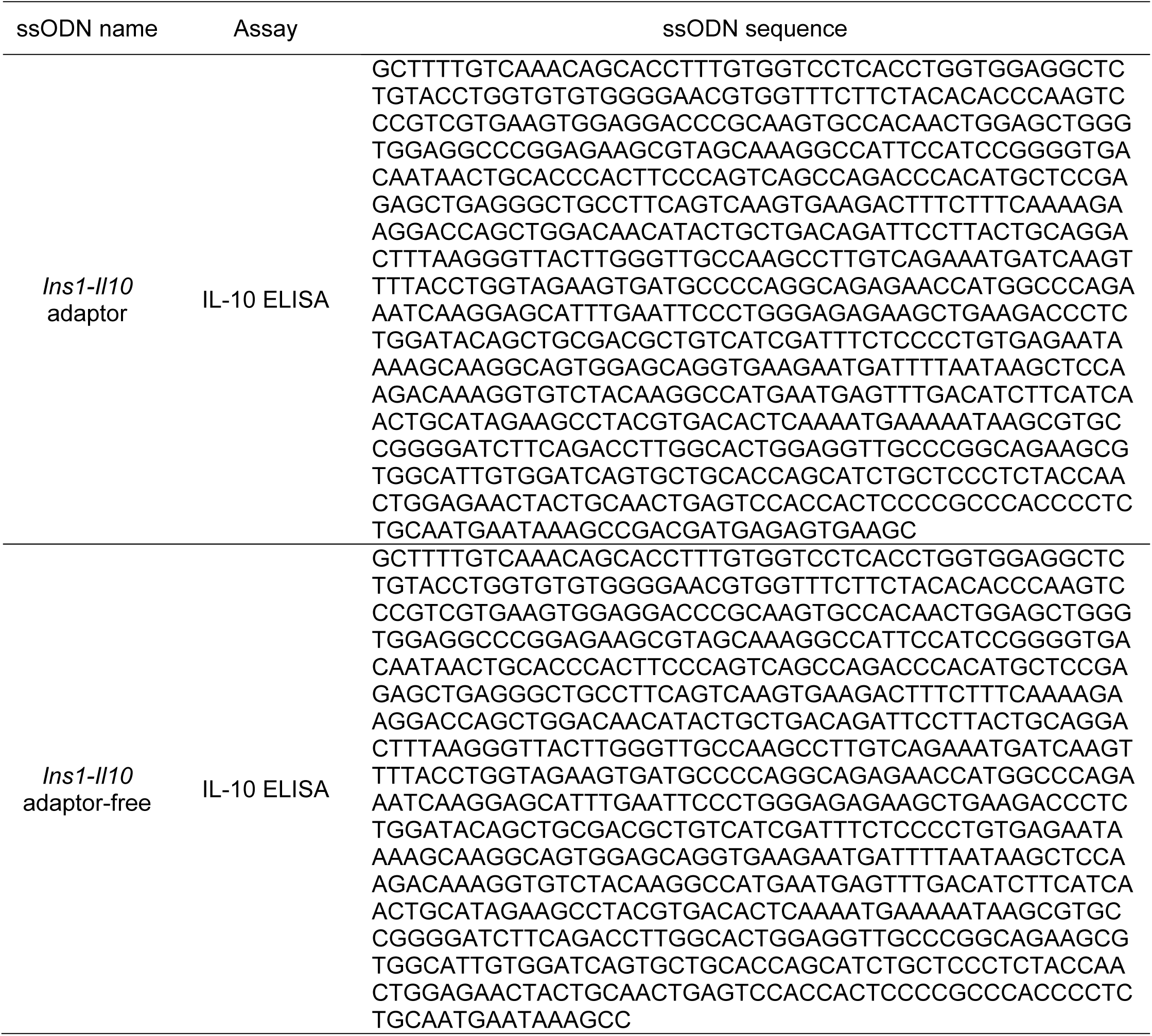
Sequences of long ssODNs for *IL-10* knock-in.

**Figure S1.**
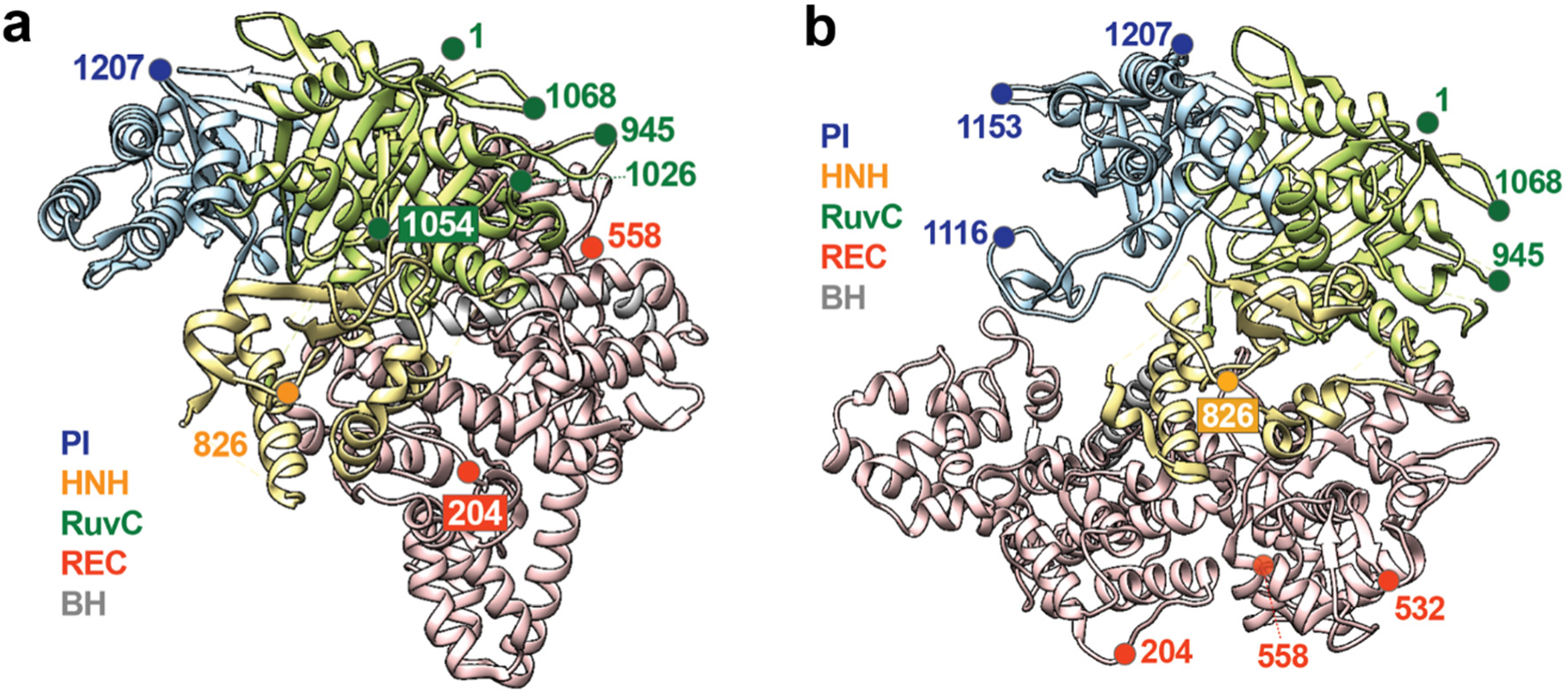
Selection of Cas9 labeling sites based on crystal structures. (a) Structure of apo-Cas9 (PDB ID: 4CMP).1 Four residues (1, 532, 1116, 1153) are not assigned at the structure possibly due to the high flexibility. We assumed that those sites are surface-exposed based on the nucleic-acid-bound structures and/or high flexibility of the loops they belong to. (b) Structure of gRNA-bound Cas9 (PDB ID: 4ZT0).2,3 Only residue 558 is projected toward the interior of the protein, indicating that labeling at this site can inhibit the formation of the correct ribonucleoprotein (RNP) structure. Cas9 exhibits a large conformational change, especially at the recognition (REC) lobe, upon gRNA binding (residues 204, 532, 558). PI domain is in blue, HNH domain is in yellow, RuvC domain is in green, REC domain is in red, and BH is in gray.

**Figure S2.**
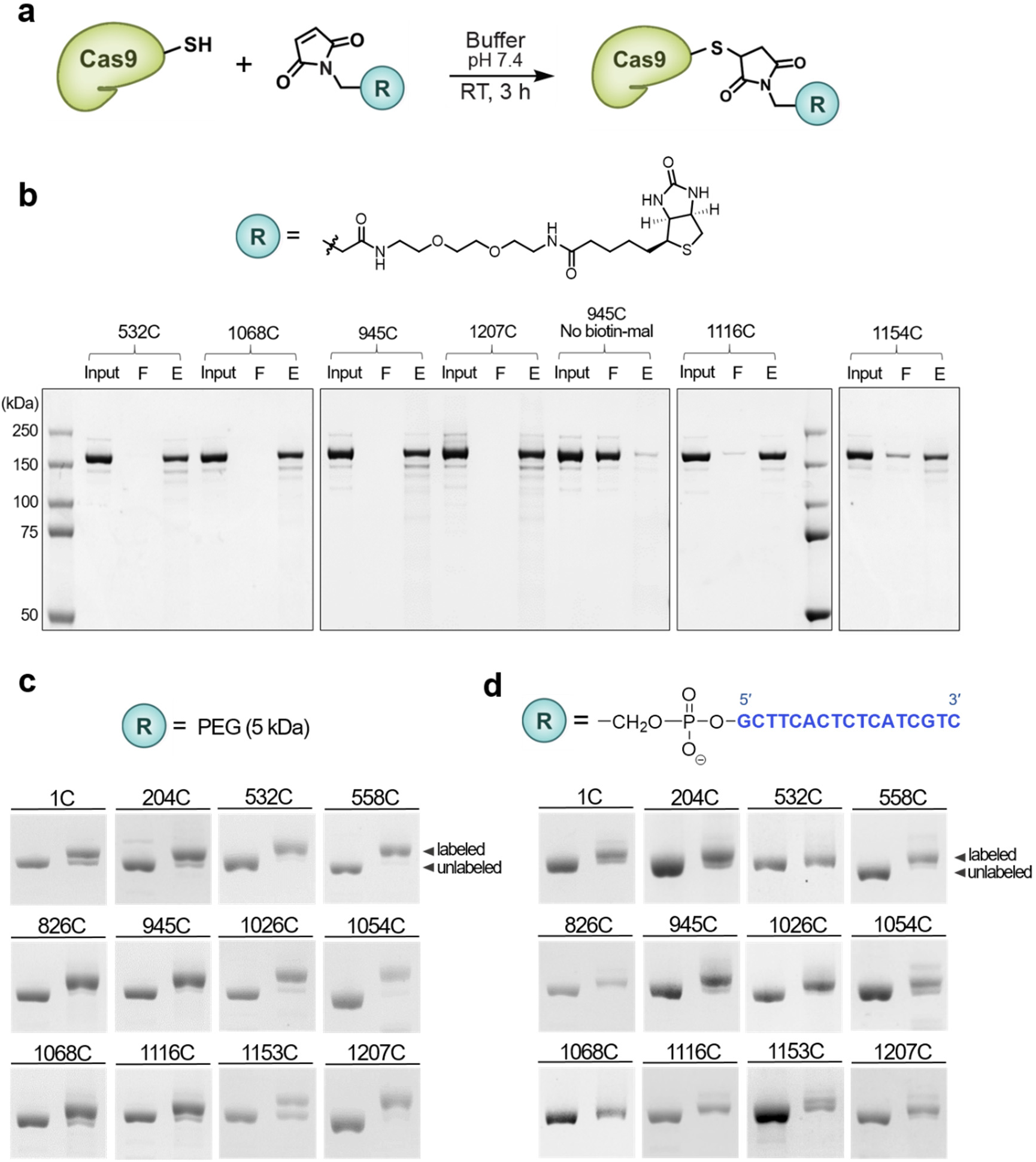
(a) Schematic of the site-specific labeling of Cas9 single-cysteine mutants by thiol-maleimide conjugation. (b) Biotin-maleimide was reacted with a cysteine on Cas9. The reaction mixture was subjected to pull-down by streptavidin beads to separate between unlabeled (flow-through, F) and biotinylated (eluate, E) Cas9. Each fraction was analyzed by SDS-PAGE followed by Coomassie staining. (c) PEG-maleimide was reacted with a cysteine on Cas9. (d) The adaptor oligonucleotide with a 5’-maleimide group was reacted with a cysteine on Cas9. The degree of labeling was monitored through SDS-PAGE followed by Coomassie staining for PEG and DNA labeling.

**Figure S3.**
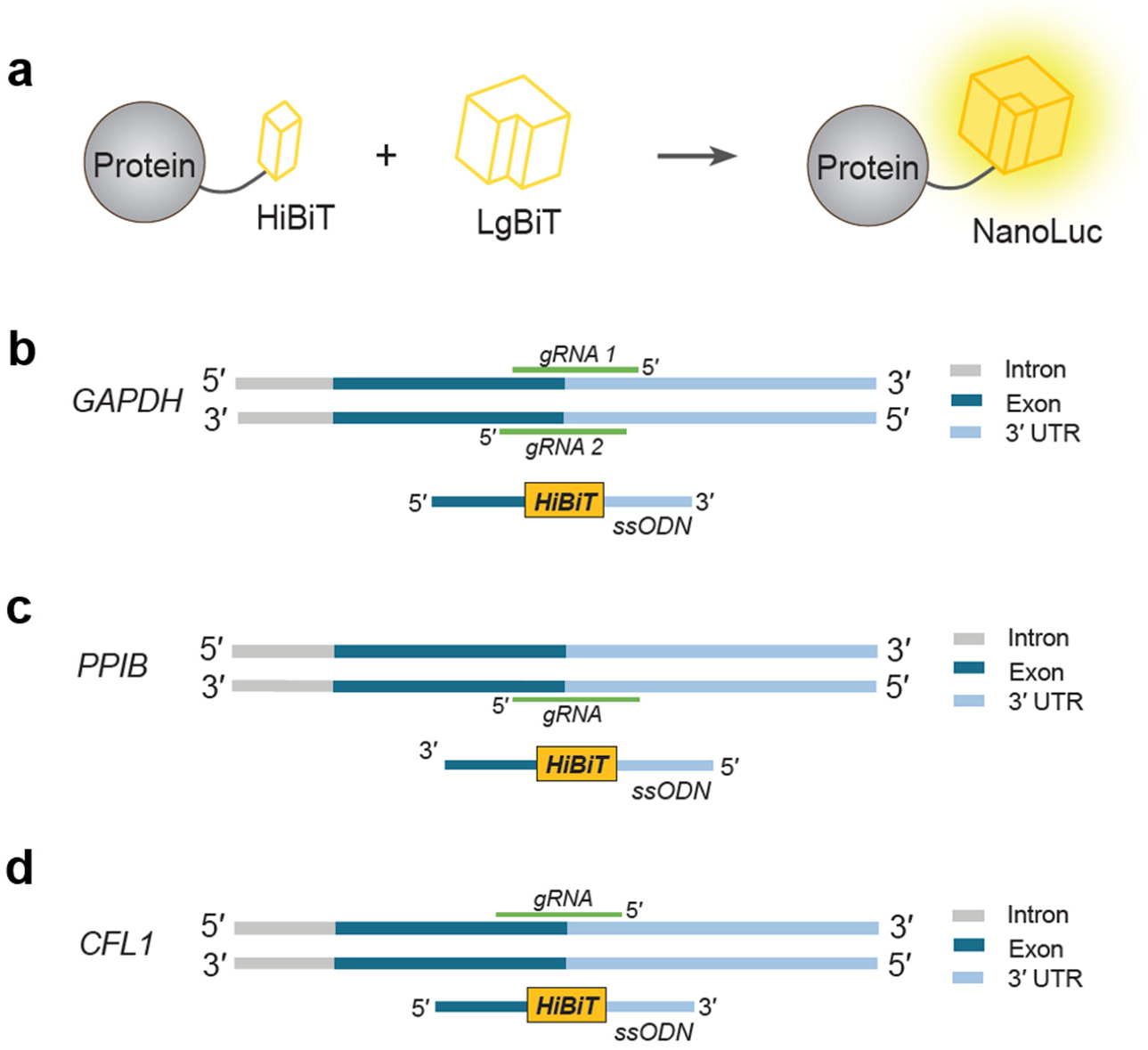
Schematic of the HiBiT assay to check the HDR-mediated knock-in of the 33-nt DNA fragment. (a) *HiBiT* sequence knock-in right before the stop codon of the gene of interest results in the expression of a fusion protein having a C-terminal HiBiT tag, which is a small fragment of the NanoLuc luciferase. When an excess amount of the other fragment of NanoLuc (LgBiT) is supplied, a fully functional NanoLuc is reconstituted. The resulting luminescence signal is proportional to the HDR efficiency. (b) Design strategy for *HiBiT* knock-in at the *GAPDH* locus. gRNA 1 was used for genome editing in Figure 2, and gRNA 2 was used in Figure 3a. (c-d) Design strategy for *HiBiT* knock-in at the *PPIB* locus (c) and the *CFL1* locus (d).

**Figure S4.**
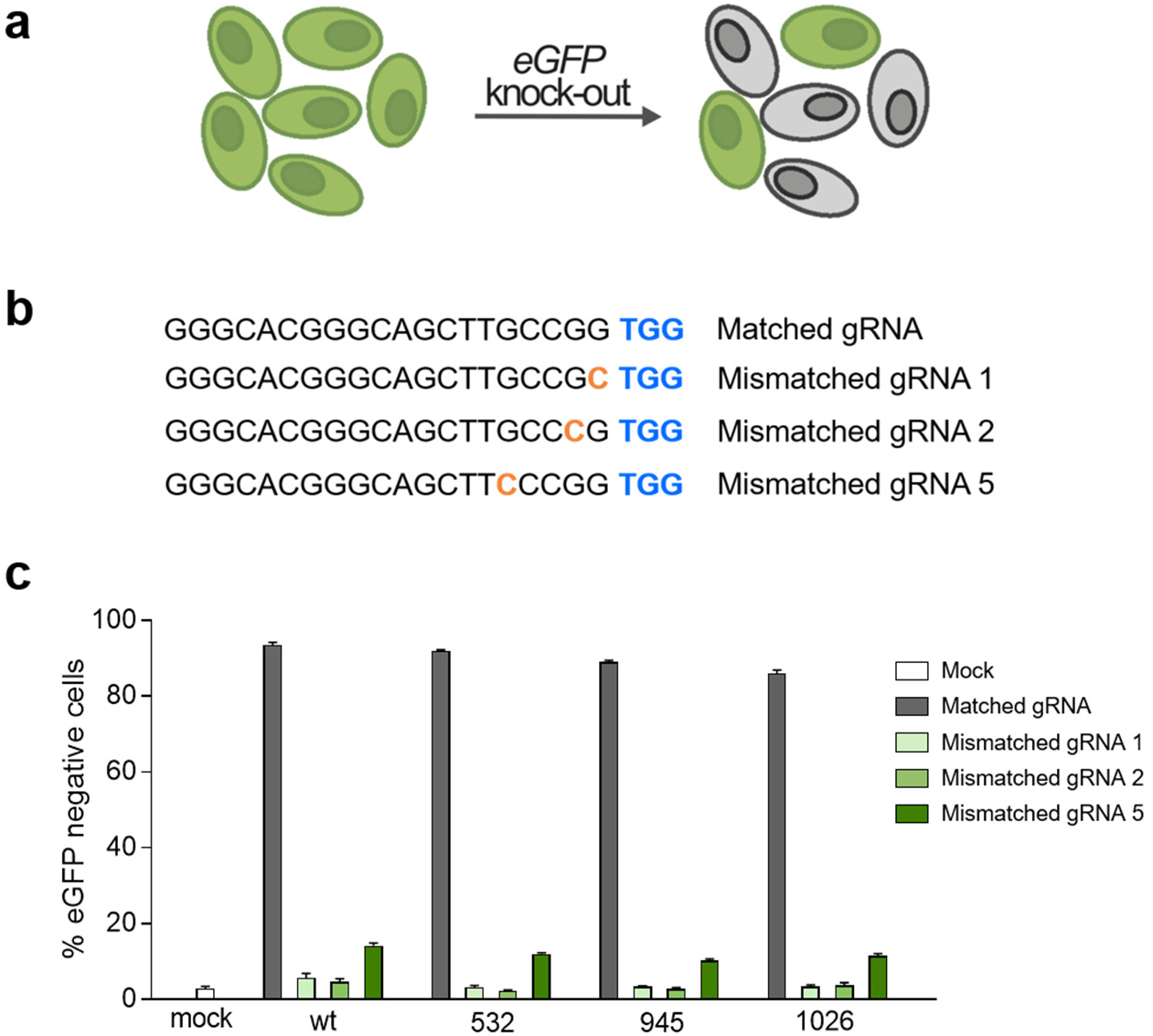
(a) Schematic of the *eGFP* knock-out assay to investigate the off-target profile of the Cas9-adaptor conjugates. The *eGFP·PEST* gene stably expressed in U2OS cells is targeted by Cas9 RNP using matched and mismatched gRNAs. (b) Sequences of the gRNAs. Mismatch sites are in orange. PAM sequences are highlighted in blue. (c) Results of the *eGFP* knock-out assay. Cells were nucleofected with 10 pmol of RNP and were incubated for 48 h followed by nuclei staining and fluorescence imaging. Unlabeled wildtype Cas9 (wt) and Cas9-adaptor labeled at the indicated residues were used. Error bars represent standard deviation from ≥ four technical replicates.

**Figure S5.**
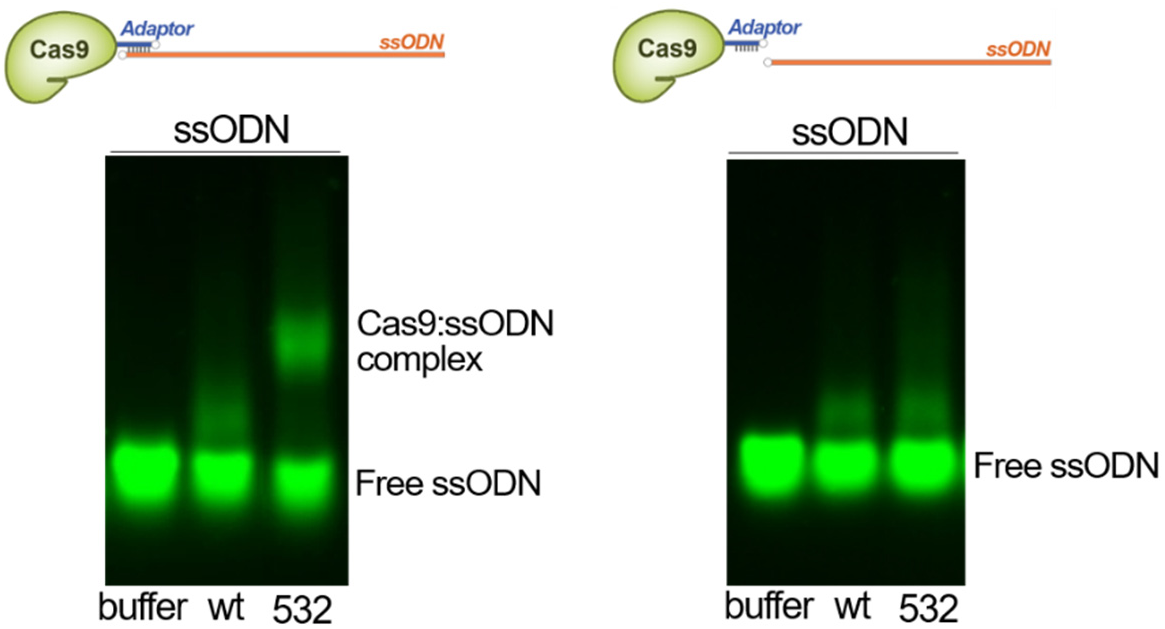
Electrophoretic mobility shift assay to check the binding between Cas9-adaptor conjugates and ssODN. When the ssODN contained the adaptor-binding sequence, the specific Cas9:ssODN complex was observed. In contrast, only non-specific binding patterns were observed when the ssODN did not have the corresponding sequence or when the unlabeled wildtype Cas9 (wt) was used. The ssODN for *HiBiT* knock-in at the *GAPDH* locus was used. Even though the lanes are not contiguous, they are all from a single gel.

**Figure S6.**
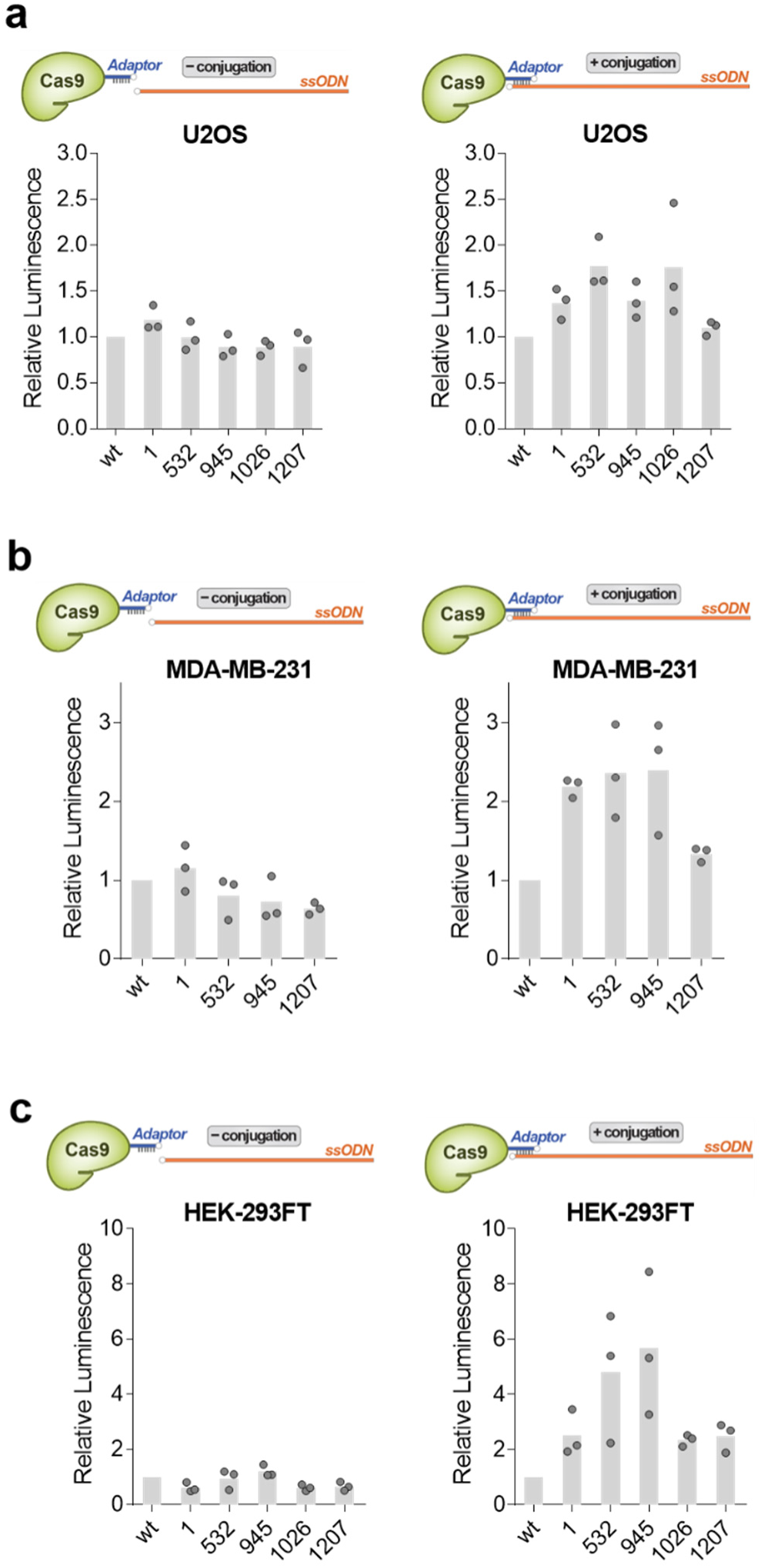
(a-c) *HiBiT* knock-in results in U2OS cells (a), HEK-293FT cells (b), MDA-MB-231 cells (c). The panels on the left show luminescence intensities using the separate Cas9/ssODN system. The panels on the right show luminescence intensities from Cas9:ssODN conjugates. Unlabeled wildtype Cas9 (wt) and Cas9-adaptors labeled at the indicated residues were used. All data from biological replicates are shown.

**Figure S7.**
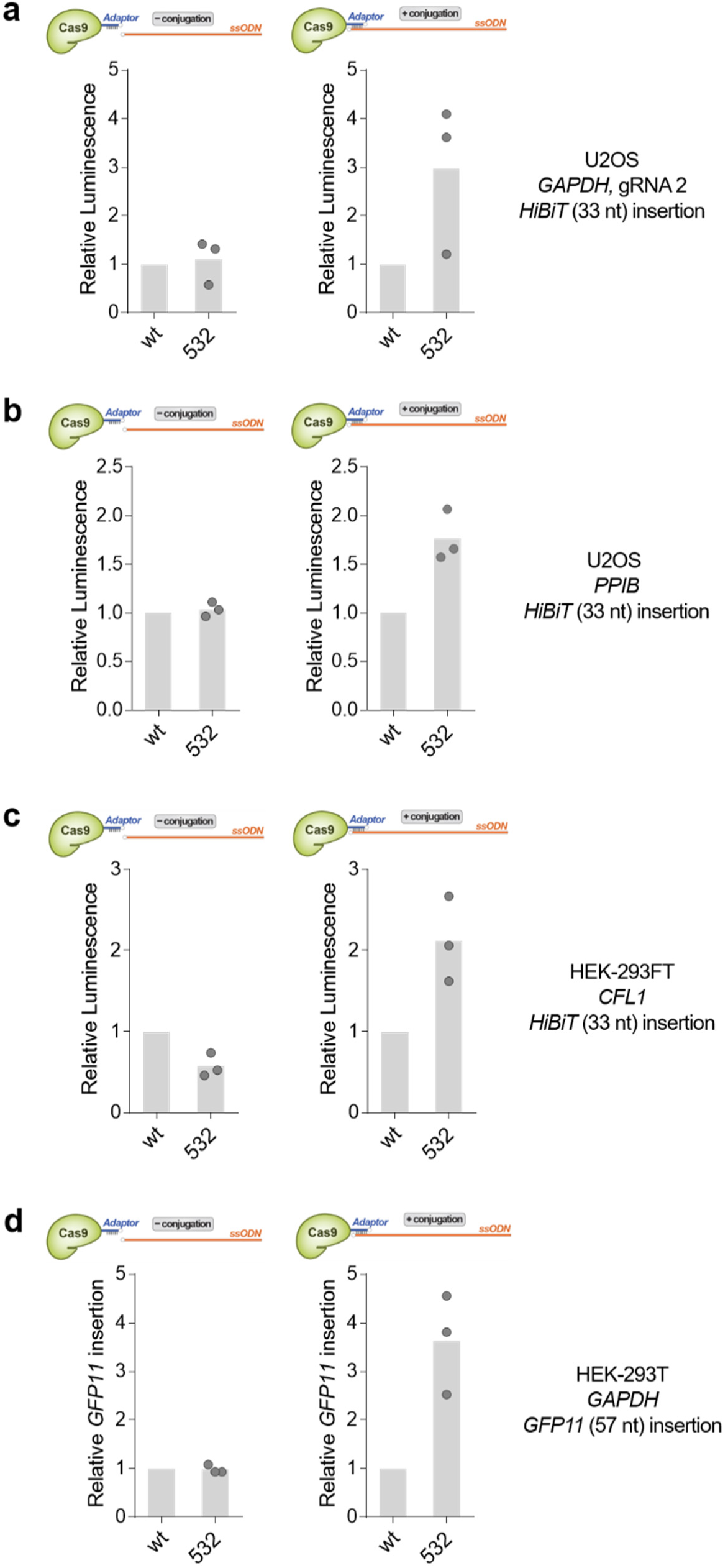
(a) Another *GADPH*-targeting gRNA was used for *HiBiT* knock-in. (b) *PPIB* locus was targeted for *HiBiT* knock-in. (c) *CFL1* locus was targeted for *HiBiT* knock-in. (d) The *GFP11* sequence was inserted at the *GAPDH* locus. Either a separate Cas9/ssODN system (left panels) or a Cas9:ssODN conjugate (right panels) was used. Unlabeled wildtype Cas9 (wt) and Cas9-adaptor labeled at residue 532 were used. All data from biological replicates are shown.

**Figure S8.**
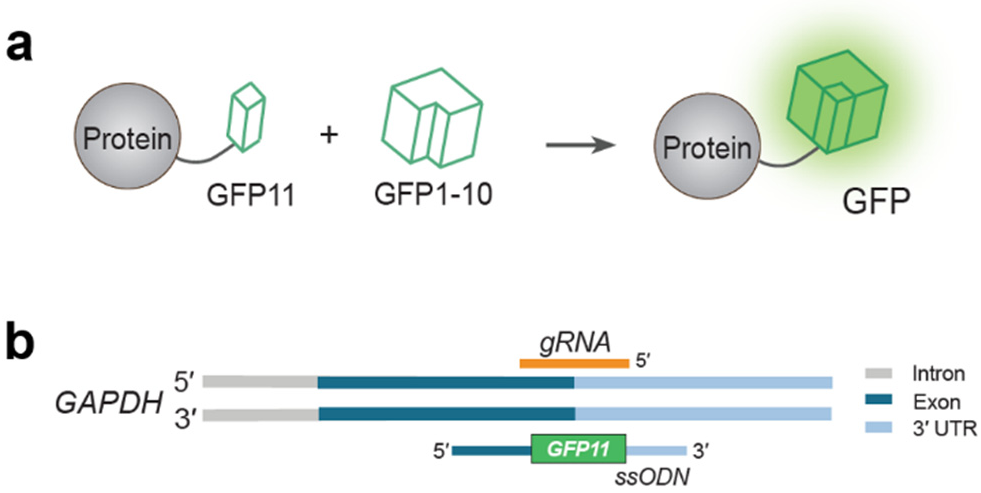
GFP complementation assay to check the HDR-mediated insertion of the 57-nt *GFP11* fragment.^4^ (a) *GFP11* sequence knock-in right before the stop codon of the gene of interest results in the expression of a fusion protein having a C-terminal GFP11 tag. When the other fragment of GFP (GFP1-10) is supplied, a fully functional GFP is reconstituted, and the fluorescence signal can be detected. (b) Design strategy for *GFP11* knock-in at the *GAPDH* locus.

**Figure S9.**
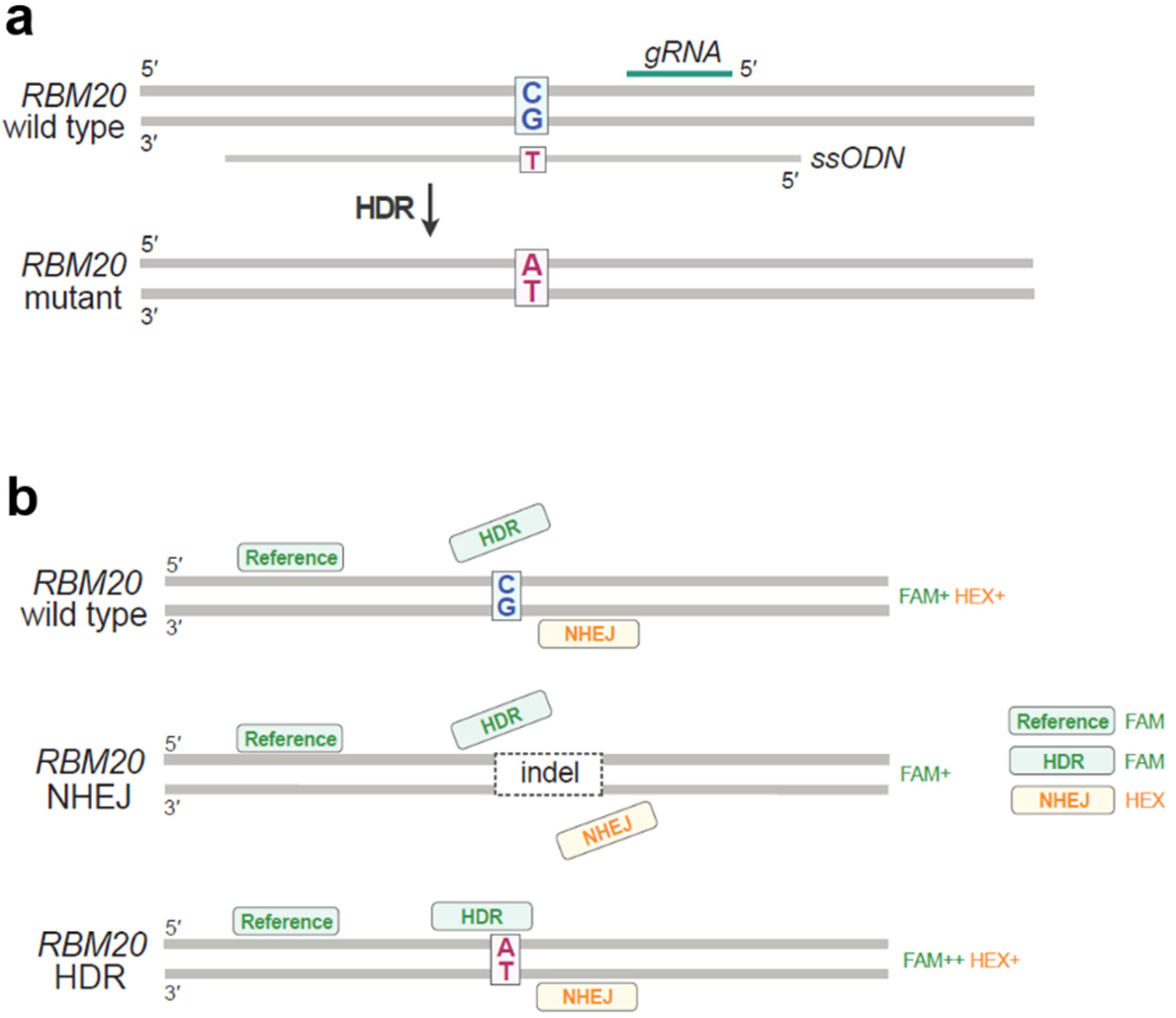
Droplet digital PCR-based quantification of single-nucleotide exchange at the *RBM20* locus. (a) One of the CG pair is replaced by AT pair. (b) Schematic of the droplet digital PCR-based quantification of NHEJ and HDR.^5,6^ The reference probe can bind to all alleles while the HDR probe binds only to the precisely edited allele. The NHEJ probe is a drop-off probe that cannot bind to the NHEJ-repaired allele. Each probe is labeled with a fluorophore-quencher pair. During the PCR, DNA-bound probes are hydrolyzed by the exonuclease activity of the DNA polymerase. Therefore, fluorophores and quenchers move apart from each other, providing fluorescence signals.

**Figure S10.**
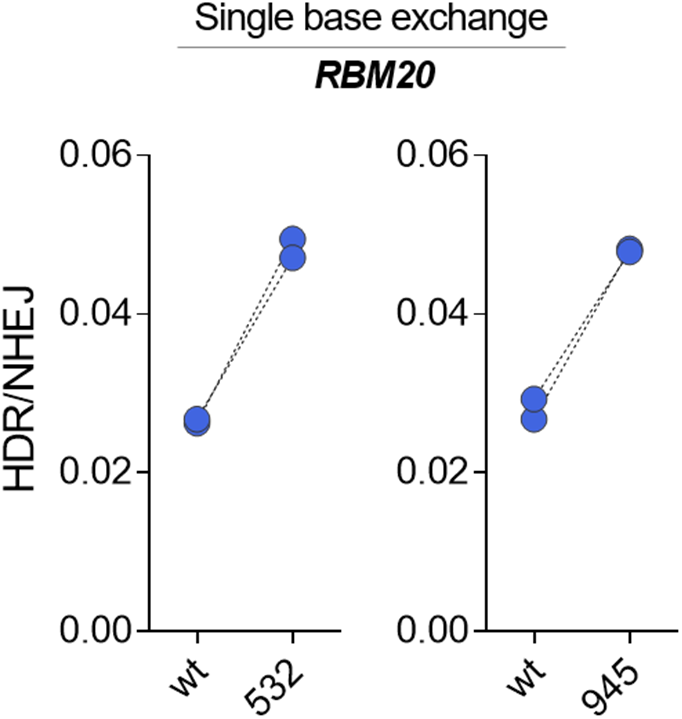
Single base exchange at the *RBM20* locus was promoted by Cas9-ssODN conjugation in HEK-293FT cells. Unlabeled wildtype Cas9 (wt) and Cas9-adaptor labeled at the indicated residues were used. Another gRNA/ssODN pair (gRNA 2 from Table S2 and ssODN 2 from Table S3) was used. All data from biological replicates are shown.

**Figure S11.**
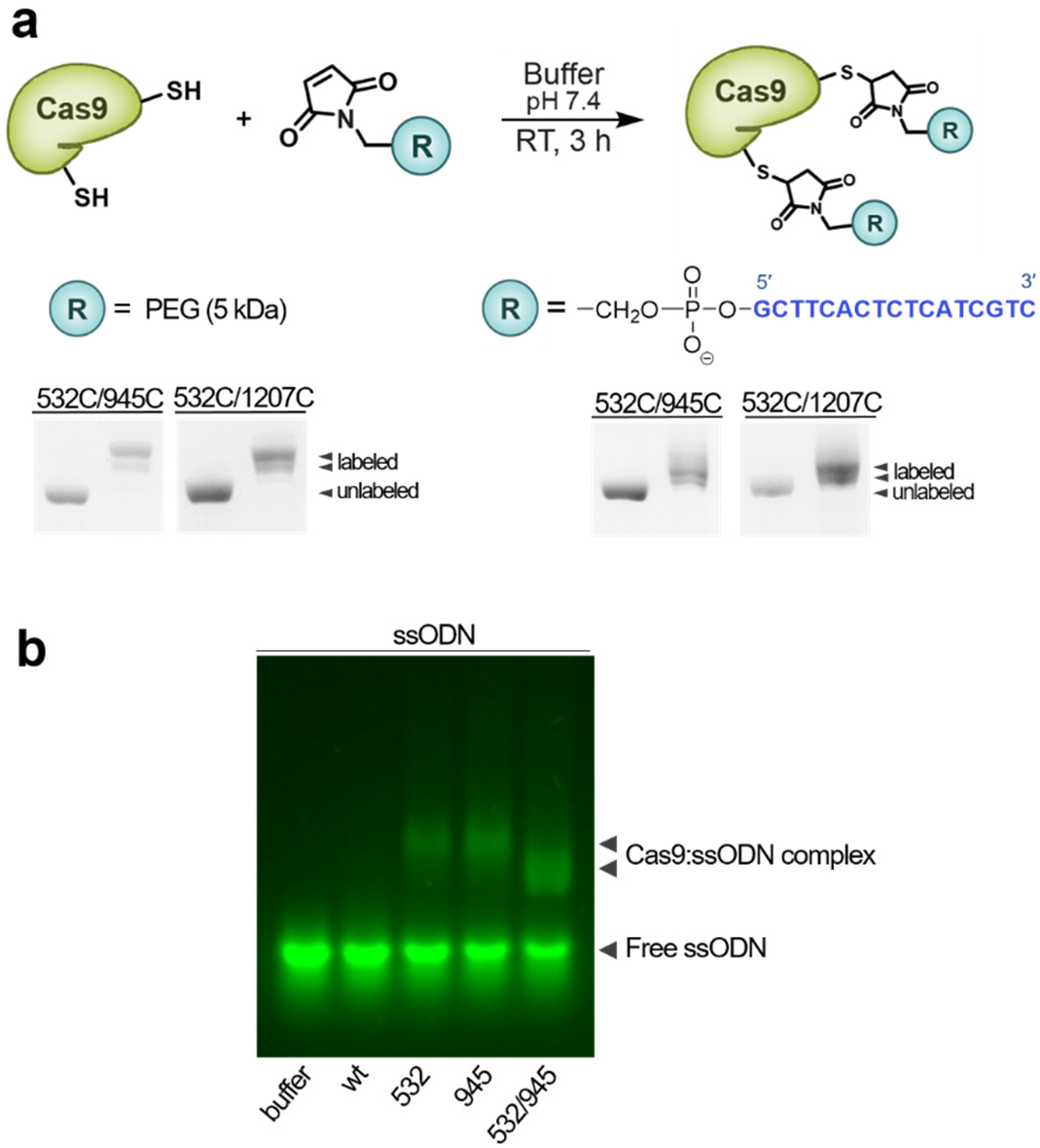
(a) Site-specific labeling of Cas9 mutants at two cysteine residues using thiol-maleimide conjugation. The degree of labeling was measured through SDS-PAGE followed by Coomassie staining. (b) An electrophoretic mobility shift assay was performed using Cas9-adaptors and the ssODN specific for *GAPDH HiBiT* tagging that contained the adaptor-binding sequence. Unlabeled wildtype Cas9 (wt) and Cas9-adaptors labeled at the indicated residues were used. The RNP and ssODN were used at a molar ratio of 1:2.

**Figure S12.**
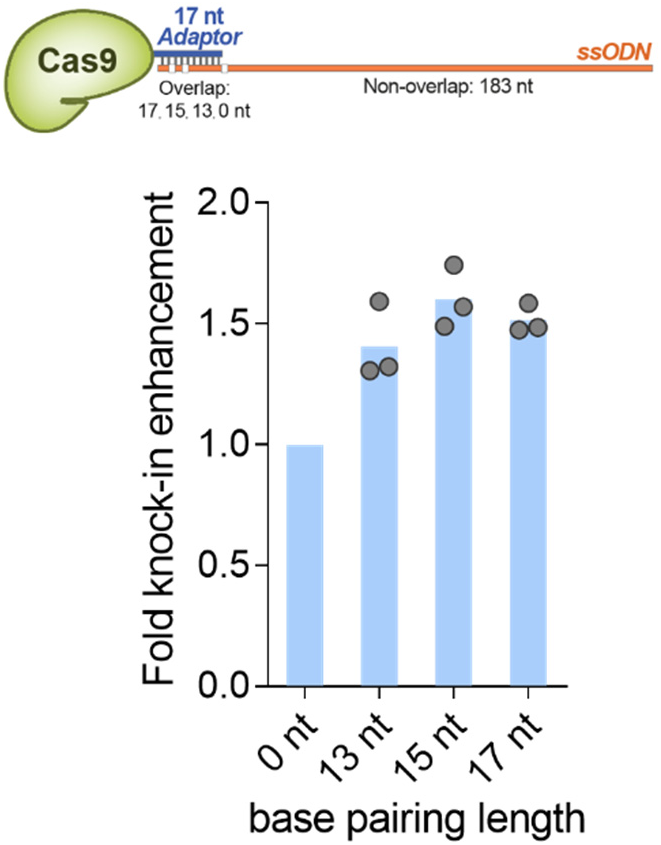
Effect of the base-pairing length on the HDR-enhancing capability of the Cas9:ssODN conjugate. *HiBiT* sequence insertion was employed as a test HDR assay in U2OS cells using the Cas9-adaptor labeled at residue 945. All data points from biological replicates are shown.

**Figure S13.**
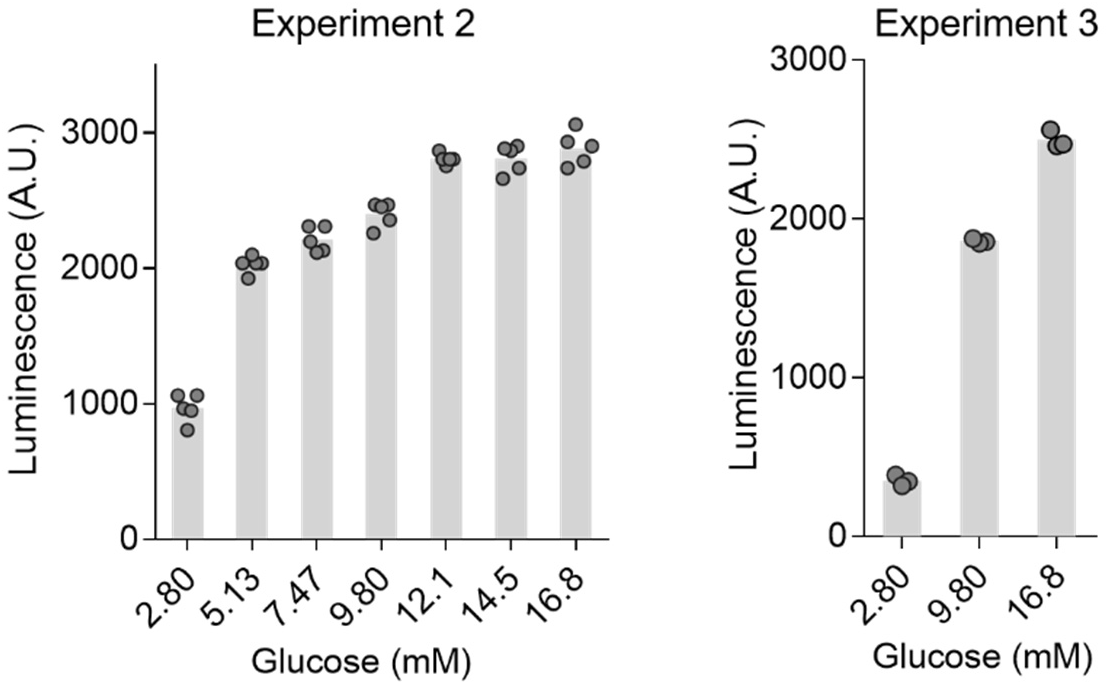
Glucose-stimulated HiBiT peptide secretion from edited INS-1E cells in independent experiments. All data points from technical replicates are shown.

**Figure S14.**
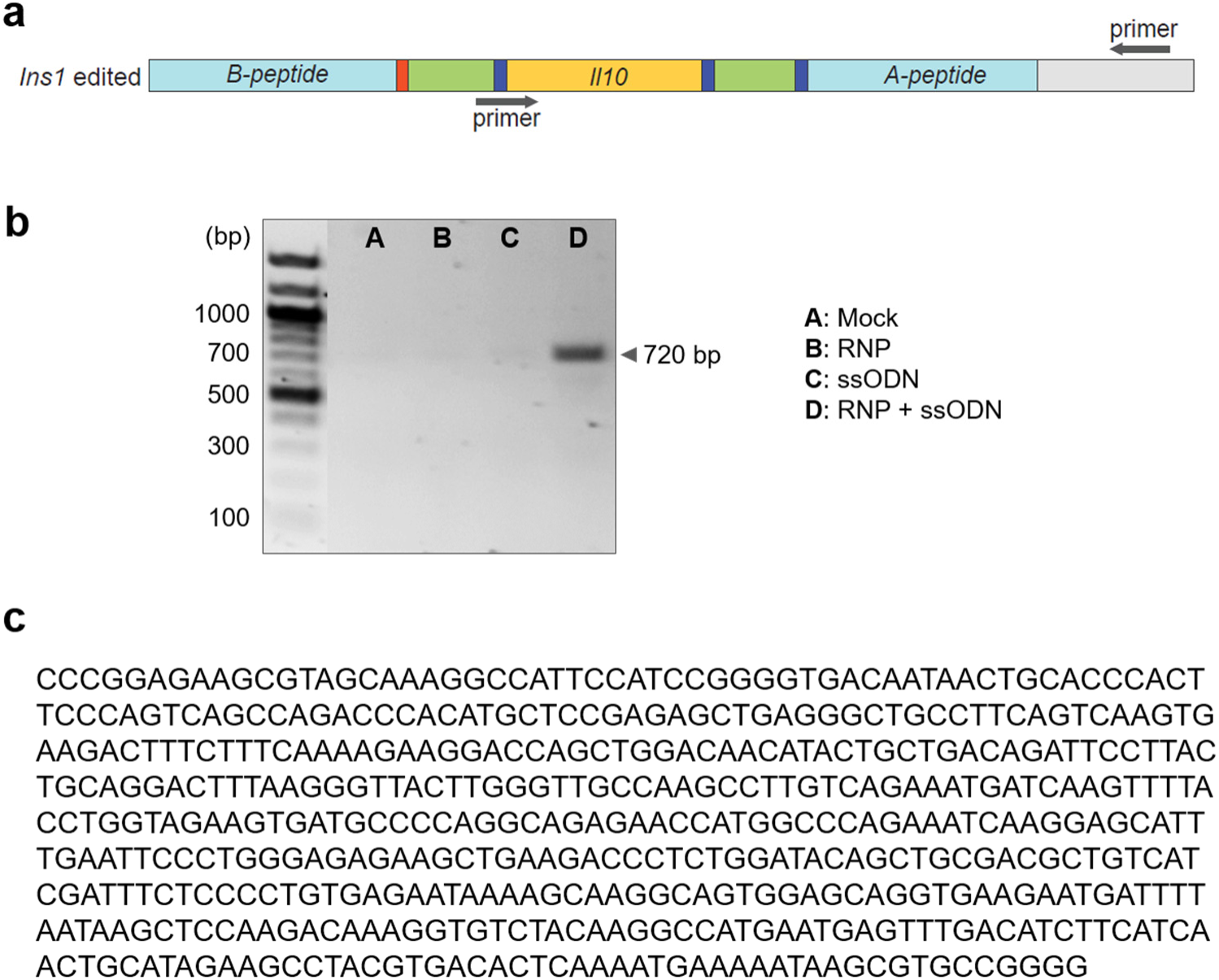
Confirmation of *Il0* knock-in by PCR. (a) Primers specific for the knock-in sequence was used. (b) Genomic DNA was extracted from cells 72 h after transfection, and PCR was performed followed by agarose gel electrophoresis and ethidium bromide staining. (c) *Il10* knock-in sequence confirmed by Sanger sequencing.

**Figure S15.**
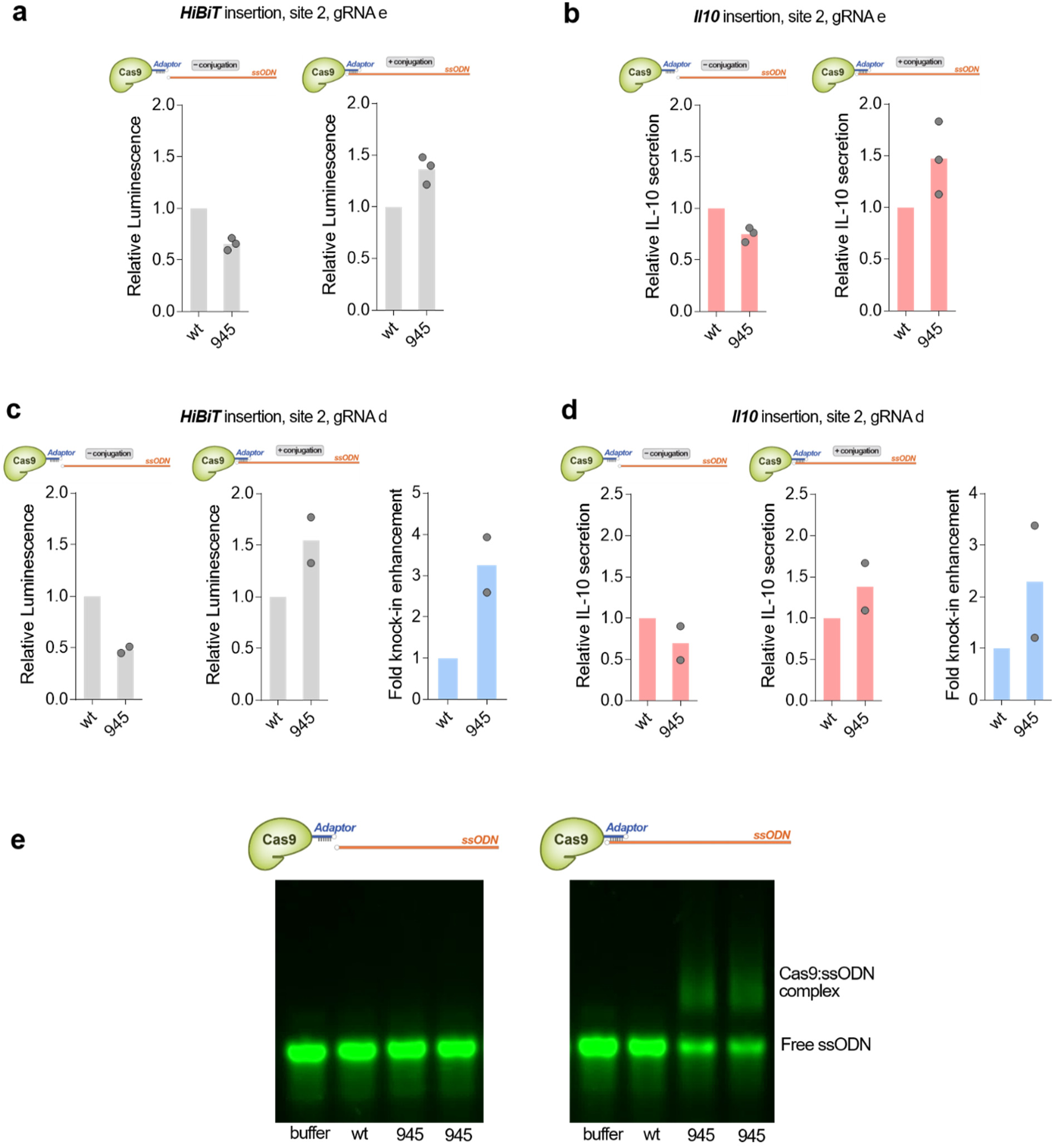
Cas9:ssODN conjugate enhanced precision genome editing in INS-1E cells. (a–d) Both *HiBiT* knock-in and *Il0* knock-in were promoted by Cas9-ssODN conjugation when two different gRNAs were tested. Unlabeled wildtype Cas9 (wt) and Cas9-adaptor labeled at residue 945 were used. All data from biological replicates are shown. (e) Electrophoretic mobility shift assay to check the binding between Cas9-adaptor and long ssODNs for *Il0* knock-in. The specific Cas9:ssODN complex was observed only when both Cas9 and ssODN contained the complementary adaptor sequences. All lanes are from a single gel. Unlabeled wildtype Cas9 (wt) and Cas9-adaptor labeled at residue 945 from different batches were used.

